# Spatial maps in piriform cortex during olfactory navigation

**DOI:** 10.1101/2020.02.18.935494

**Authors:** Cindy Poo, Gautam Agarwal, Niccolò Bonacchi, Zachary Mainen

## Abstract

Odors are a fundamental part of the sensory environment used by animals to inform behaviors such as foraging and navigation^1, 2^. Primary olfactory (piriform) cortex is thought to be dedicated to encoding odor identity^3–8^. Here, using neural ensemble recordings in freely moving rats performing a novel odor-cued spatial choice task, we show that posterior piriform cortex neurons also carry a robust spatial map of the environment. Piriform spatial maps were stable across behavioral contexts independent of olfactory drive or reward availability, and the accuracy of spatial information carried by individual neurons depended on the strength of their functional coupling to the hippocampal theta rhythm. Ensembles of piriform neurons concurrently represented odor identity as well as spatial locations of animals, forming an “olfactory-place map”. Our results reveal a previously unknown function for piriform cortex in spatial cognition and suggest that it is well-suited to form odor-place associations and guide olfactory cued spatial navigation.

## Introduction

Olfaction is critical to animals navigating their environments for valuable resources^1^. Animals instinctively use odor memories to guide spatial choices^9, 10^ and odors are widely used in the study of spatial memory and navigation^11^. Cortical structures for odor perception and spatial memory are evolutionarily and developmentally linked, together forming an allocortex (consisting of olfactory, hippocampal, and entorhinal cortices). Olfaction and spatial memory systems are therefore intimately related as reflected by animal behavior, evolution, and circuit anatomy. Moreover, while the olfactory (piriform) cortex receives direct sensory input via olfactory bulb projection neurons, its circuit architecture shares striking resemblances to the hippocampus, with ongoing plasticity of broadly distributed and unstructured recurrent connections^12–16^. This has prompted conjectures that these structures implement similar associative learning functions^13, 17^. Despite these suggestive parallels, experimental evidence has been largely consistent with a distinct functional division between piriform and hippocampal systems in attributing odor encoding to piriform, and spatial navigation and memory to the hippocampal network^3, 5–8, 18, 19^. However, if spatial information was present in the piriform cortex (PCx) it would be well-positioned to function as part of the spatial memory system.

Within the PCx, anatomical and neurophysiological studies have reported differences between anterior and posterior regions (aPCx and pPCx)^16, 20–26^. While both regions are characterized by recurrent circuitry, aPCx receives more inputs from the olfactory bulb and other olfactory regions, whereas pPCx is more strongly connected to higher-order association and cognitive regions^16, 20–24, 27–29^. The primary function of aPCx has generally been thought of as being dedicated to odor identification and discrimination^3, 30–32^, while much less is known about the functions of pPCx^27, 33^.

### Odor-cued Spatial Choice Task

We developed a novel odor-cued spatial choice task that challenged rats to use spatial information along with odor identity to navigate to a specific location for reward. We recorded from pPCx and area CA1 of the hippocampus in rats performing this task (**Fig. 1**, **Extended Data Fig. 1**). Our behavioral arena consisted of an elevated plus-maze with a one square meter footprint that was positioned in a room with visual landmarks. At the end of each arm was a port, each of which could deliver either an odor cue or a water reward. Trials began when an LED was activated at one of the four ports. Rats learned to insert their snouts into the lighted port in order to receive one of four odors chosen at random (**Fig. 1a and supplementary video**). Odors were transient and only available when rats held their snout within the sampling port (**Fig. 1b**). Following odor sampling, rats had to use the odor identity to select the correct spatial goal associated with the odor and to navigate there in order to collect water reward (**Fig 1c**, **Extended Data Fig. 2**).

**Figure 1.**
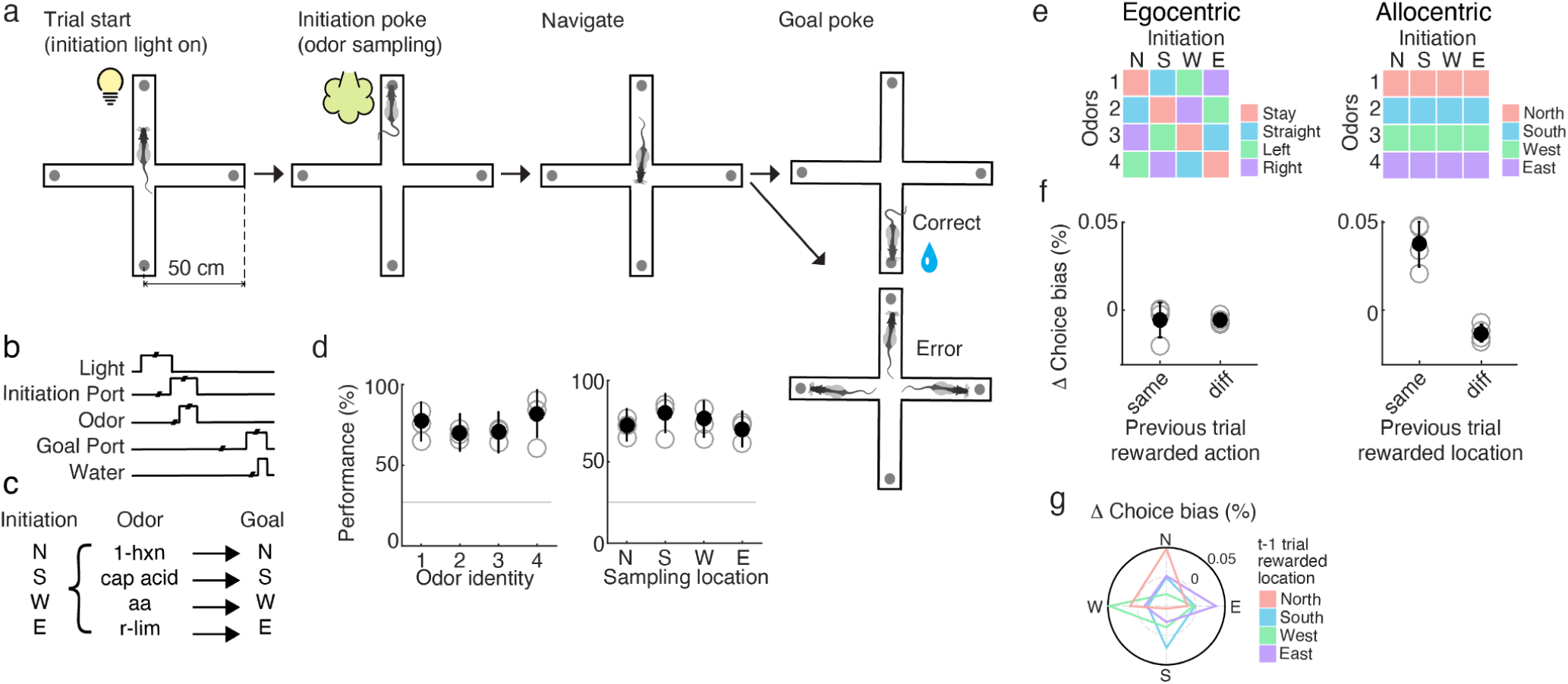
Odor-cued allocentric spatial choice task. (**a**) Task structure and behavioral arena. Rats behave on a 1m x 1m elevated plus-maze with a nose port at the end of each arm. Each port can deliver either a light cue, odor, or water reward. The rat initiates a trial by following a light cue to poke into a nose port to receive an odor (1 out of 4). An odor can be presented at any one of the four possible initiation ports. The rat must then choose one of the four possible ports (North, South, West, or East port) to retrieve a water reward. Each odor was associated with a water reward at a fixed goal location. (**b**) Schematic of temporal events in trials. (**c**) Odor to reward location mapping. Each odor can be sampled at any initiation port, but odor identity determined the spatial location of where water reward was available. (**d**) Behavioral performance of rats was similar across all odor identities and initiation (odor sampling) locations.(**e**) Schematic illustrating the allocentric (**left**) or egocentric (**right**) rule that must be followed by rats in order to correctly perform the task given all possible odor and sampling locations. (**f**) Reward for a particular action does not bias the choice probability for the same action in the following trial (**left**). Reward at a particular location biases the choice probability of rats towards the same location in the following trial (**right**). (**g**) Polar plot shows that given reward at specific port locations, rats were positively biased towards choosing the same location on subsequent trials (see Full Methods).

Critically, trial initiation cues were randomly interleaved across the maze arms so that odors were presented with equal frequency at all four ports. Therefore, adopting a location-based (allocentric) strategy would be simpler than the alternative of learning and applying a different direction-based (egocentric) action rule for each odor depending on where it was sampled^34, 35^(**Fig 1e**). After training (criterion: > 70% accuracy, reached after 3-4 weeks, chance level = 25%, > 200 trials/session, see Methods), performance of rats was similar across odors, choice locations and choice directions (**Fig. 1d**).

This task differs in an important way from most rodent sensory-cued decision tasks. Most other olfactory tasks that use spatial responses (e.g. left vs. right choice) do not require use of a spatial map because they can be solved by directly associating an odor cue with an action. A signature of such decision tasks is that animals show a win-stay bias, repeating a successful action following reward^36^. In the odor-cued spatial choice task here, rats instead show a bias to return to the same location following a successful trial, but no bias to perform the same action **(Fig. 1f and g**, 6 rats, ∼40,000 trials, see Full Methods). In other words, the majority of errors are made when rats disregard current trial odor cue, in order to revisit the most recently rewarded location. This suggests that rats are indeed using spatial maps in conjunction with olfaction in this task. Interestingly, animals performed with higher accuracy when the odor-cued goal location is congruent with odor sampling location (‘stay’ trial type, **Extended Data Fig. 2c**), and errors that were caused by animals staying at the odor sampling port were less common than other actions (**Extended Data Fig. 2d**).

### Spatial locations are encoded in piriform cortex

We recorded from pPCx during the odor-cued spatial navigation task using chronically implanted tetrodes, and found that neurons fired differently depending on odor identity as well as on the location of the rat (**Fig. 2a-d**). Individual pPCx neurons displayed a range of odor and location selectivity, from strictly spatial (**Fig. 2a**) to purely odor selective (**Fig. 2b**). These selectivity properties did not correlate with extracellular electrophysiological signatures associated with specific cell types such as spike waveform or overall firing rate (data not shown). To account for the possibility that the spatially-restricted firing of pPCx neurons reflected the influence of stray odors on the maze (potentially deposited by the rat)^37^, each recording session was split into two blocks, separated by a 15-20 min interval during which the rat was removed and the maze and ports were thoroughly cleaned with disinfectant and an enzymatic cleaner.

**Figure 2.**
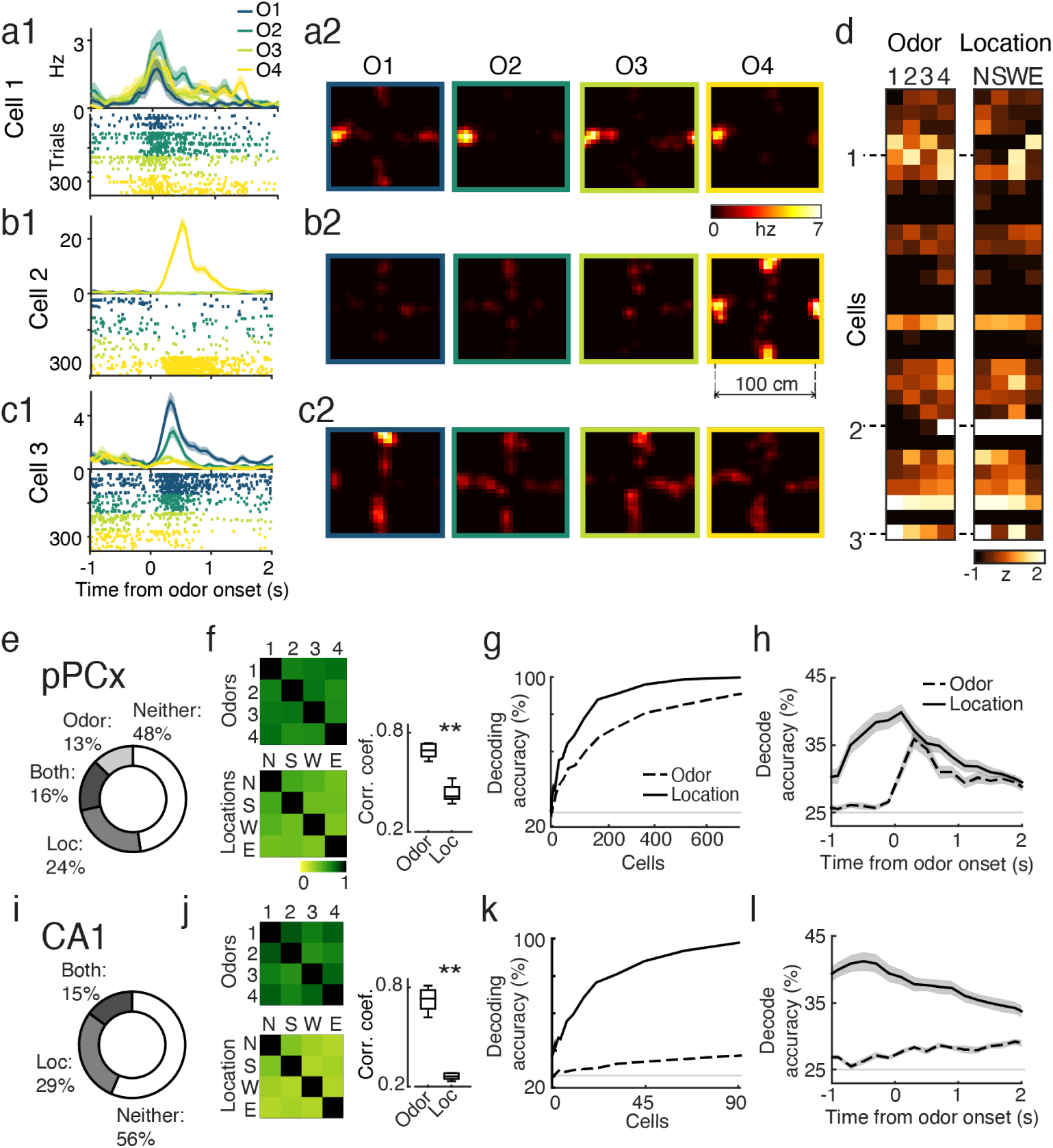
Spatial representations in the piriform cortex. (**a-d**) One example recording session of simultaneously recorded pPCx population (n = 30 cells). (**a1, b1, c1**) Perievent time histogram (PETH, top) and raster plots (bottom) reveal odor selectivity (odors labeled O1-4). Spike times were aligned to odor onset. (**a2, b2, c2**) Firing rate heat map for different odors (O1-4) normalized for occupancy on maze. Heat maps were generated by concatenating all trials for an individual odor from -1 to 2 s around initiation port entry (similar time period as shown in raster and PETH, see Full Methods) (**d**) Z-scored firing rate of neurons show that neurons were differentially activated for odors and locations. Mean z-scored firing rate for odors was taken between 0-1 s after odor onset. Mean z-scored firing rate for locations was calculated from occupancy normalized firing rate heat map, using 20 cm x 20 cm position bins centered around port locations. (**e**) Selectivity of pPCx neurons for odors, locations, or both. (**f**, left top) Population correlation similarity matrix obtained by computing the pairwise Pearson’s correlation coefficients for pPCx population response vectors across 4 odors. Squares along the diagonal band are 1 representing autocorrelation. (**f**, left bottom) Same analysis as left but for port locations. (**f**, right) pPCx population correlation coefficients for odor and locations shown in similarity matrices, excluding autocorrelation coefficients represented by diagonal band. r_ppc_odor = 0.69 ± 0.04, r_ppc_loc = 0.43 ± 0.05. (Wilcoxon rank-sum test, p < 0.001). Box plot central mark: median, box edges: 25th, 75th percentiles, whiskers: range of data points). (**g**) Linear classifiers decode accuracy for odor identity (dashed) and locations (solid). Lines are means of 10 subsampled pPCx pseudo-populations. (**h**) Decoding accuracy for location (solid) and odor identity (dashed) for simultaneously recorded pPCx populations (Mean ± S.E.M., n = 33 sessions; 30 ± 13 neurons/session). (**i-k**) Similar analysis as in (e-g) for CA1 population (n = 154 cells). (**j**, right) CA1 population correlation coefficients for odor and locations show in similarity matrices, excluding autocorrelation coefficients represented by diagonal band. r_ca1_odor = 0.72 ± 0.03, r_ca1_loc = 0.26 ± 0.01. (Wilcoxon rank-sum test, p < 0.001). (**l**) Decoding accuracy for location (solid) and odor identity (dashed) for simultaneously recorded CA1 populations, (Mean ± S.E.M., n = 15 sessions, 8 ± 1 cells/session).

Spatial firing rates of individual neurons across the two blocks within a session were stable across locations (**Extended Data Fig. 3,** see Full Methods). We quantified the stability of location representation between the two blocks by calculating the pairwise Pearson’s correlation coefficient for population activity across different locations in these two blocks. Values along the diagonal band in the correlation coefficient matrix represent the similarity of the pPCx population response to the same location in two recording blocks. The correlation coefficient values were significantly higher along the diagonal than off-diagonal (r_diag = 0.80 ± 0.05, r_offdiag = 0.54 ± 0.09, Wilcoxon rank sum test, p < 0.01), indicating that location representations were stable between blocks. Similar results were obtained when we examined all recorded pPCx neurons (**Extended Data Fig. 3f**, bottom. r_diag = 0.81 ± 0.02, r_offdiag = 0.47 ± 0.02, Wilcoxon rank sum test, p < 0.001). We conclude from these data that uncontrolled odors were unlikely to account for the pPCx location representations observed.

Across the entire set of pPCx recordings, we found, surprisingly, that more pPCx neurons were selective for location than for odor identity (40% vs. 29%, **Fig. 2e**). We did not see systematic biases in behavior or neural responses that might result from the non-uniform availability of odors across the maze (**Fig. 1d, Extended Data Fig. 4b-d** show a similar fraction of neurons responsive across odor identity and port locations). Overall, mean firing rates of pPCx neurons were low^33^ (**Extended Data Fig. 4a top**). There were no differences in firing rates between the subpopulations of neurons that had odor, location, or joint selectivity (**Extended Data Fig. 4a bottom, b**). Sparseness of odor responses, which is a measurement of unevenness of neural responses, was somewhat lower (less sparse) than those reported previously in aPCx^3, 30^ (**Extended Data Fig. 4e**, population sparseness = 0.36 ± 0.09; lifetime sparseness = 0.21 ± 0.05, mean ± S.D., n = 33 sessions). Correlation analysis of population response vectors showed that the pPCx population encodes spatial locations more distinctly than odors (**Fig. 2f**). There was no systematic relationship between how odors and locations were represented across the population. For example, population response vectors were not more similar between odor 1 and the north port (**Extended Data Fig. 5**). We quantified the spatial representations observed in pPCx populations by using linear classifiers to understand how well a naive downstream observer could decode odor identity or spatial location of the animal by observing firing rates of pPCx neurons (L2- regularized logistic regression with 5-fold cross-validation, see Full Methods). Pooling all recorded neurons across sessions, near perfect decoding of location (90%) could be achieved using approximately 250 pPCx neurons, whereas more than 600 neurons were required for maximum classification accuracy for odors (accuracy = 89% for 725 neurons) (**Fig. 2g**). Similar odor decoding results were obtained when pPCx spike times were aligned to first inhalation after odor onset (**Extended Data Fig. 6b**, Full Methods). Linear classifiers trained using only simultaneously recorded pPCx neurons were able to decode odor identity and spatial locations across time, showing that both odors and spatial locations are concurrently represented in piriform (**Fig. 2h**).

These results indicate that a greater fraction of pPCx neurons are involved in representing spatial location compared to odor identity and that many more pPCx neurons are required to reach a given level of odor decoding accuracy compared to what has been observed for aPCx using similar methods^3, 30–32^. This suggests that, at least in the context of our task, odor coding in pPCx could be considered fairly poor, consistent with a previous report^33^. It is worth noting that in these methods of population decoding, the decoder randomly samples neurons in the population. If, instead one uses a decoder which considers the most informative neurons first (using sparse regularization, see Full Methods), then around 150 pPCx neurons can reach ∼ 90% decoding accuracy (**Extended Data Fig. 6c, yellow line**). Therefore, our results should not be taken to exclude the possibility that a downstream area that was listening to the right subset of pPCx neurons could extract significant odor information.

### Piriform spatial maps are stable and robust

The time course of location decoding accuracy in pPCx often began well ahead of odor delivery, suggesting that spatial coding in pPCx does not depend on olfactory drive (**Fig. 2h, Extended Data Fig. 7**). However, the prominence of spatial selectivity around the odor ports also suggests the possibility that it might reflect task-related odor-predictive signals^28^. In this case, we would expect them to be specific to this epoch of the task. Alternatively, spatial information could be something akin to a cognitive map of space^34, 38^, in which case, we would expect to see representation of a spatial environment that is consistent across task epochs. To examine this, we compared the spatial selectivity of neurons around the two different kinds of task-related nose poke events, the initiation of the trial, when an odor is anticipated and the choice of goal, when a reward is anticipated. . Consistent with the ‘cognitive map’ hypothesis, spatial properties of individual neurons remained constant across these two different task epochs (**Fig. 3a-e**). Similarity matrices for population response vectors demonstrated the stability of pPCx location representation, as reflected by the diagonal axis of higher correlation coefficients (**Fig. 3b**). Indeed, a linear classifier trained using population activity during the initiation epoch was able to successfully decode location from neural activity during the reward epoch (**Fig. 3c**). This result held true even for neural activity during the inter-trial interval (ITI) when rats were not actively engaged in the task. Moreover, location representation between correct and error trials were also consistent: a linear classifier trained using neural activity in only correct trials was able to successfully decode location from error trials (**Fig. 3f, Extended Data Fig. 8**). Together, these data suggest that pPCx carries a spatial map of the environment that is relatively stable and persistent across time.

**Figure 3.**
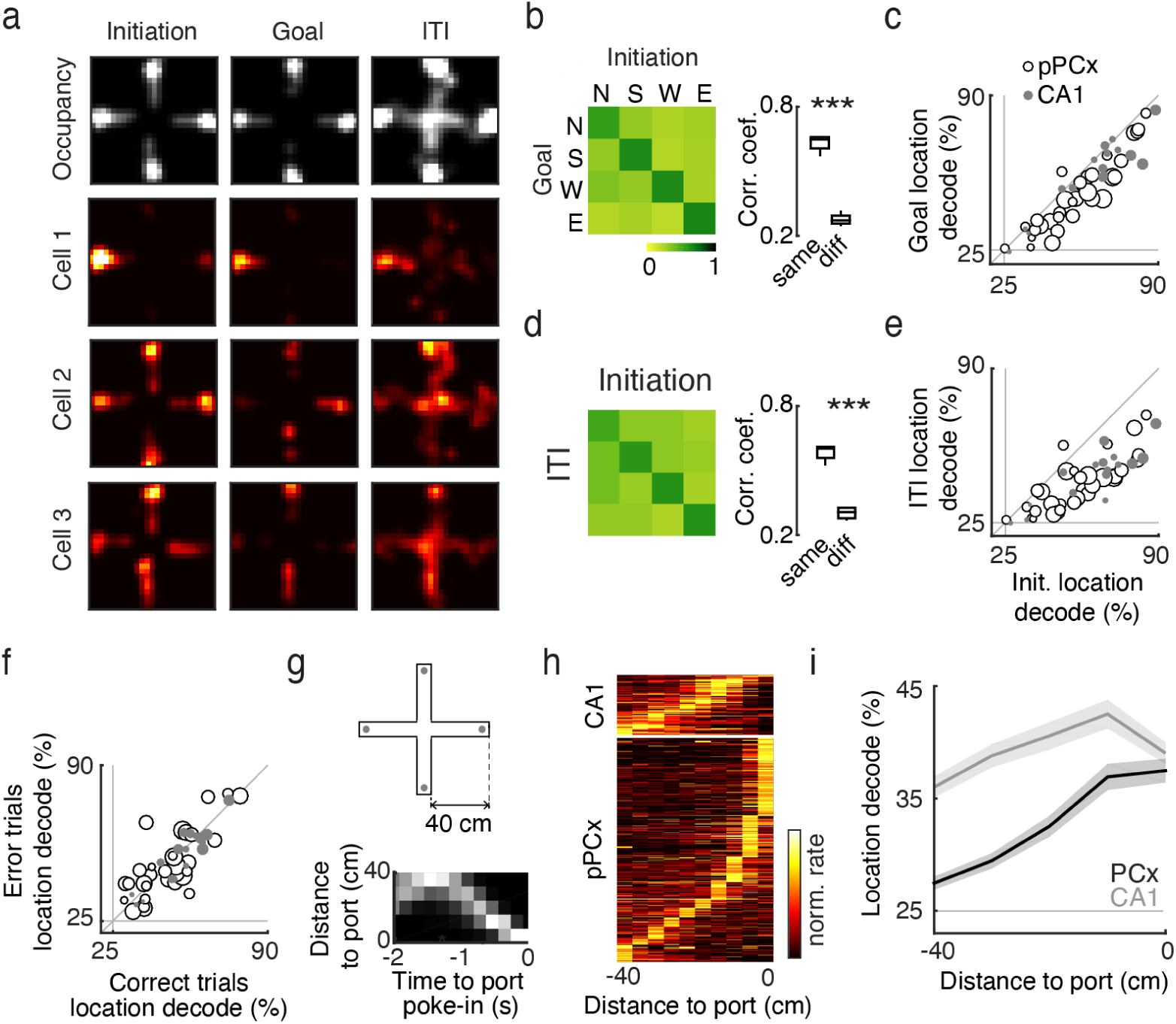
Spatial representations are robust and stable across task events. (**a**) Occupancy of rat on maze and firing rate heat maps during three behavioral epochs in an example session. All trials within a session are concatenated. Initiation epoch is -0.5 to 0.5 s around initiation port entry. Goal epoch is -0.5 to 0.5 s around goal port entry. Inter-trial-interval (ITI) starts 1.0 s after goal port entry, and has a duration of 4-6 s (randomly drawn from a uniform distribution). (**b**) Similarity matrices for spatial representation between initiation and goal epochs. Pearson’s correlation was performed on population response vectors at port locations for all location selective neurons. n = 395 pPCx cells. Correlation coefficients at the same versus different refer to diagonal versus off-diagonal values in similarity matrix. r_ppc_same = 0.63 ± 0.04, r_ppc_diff = 0.28 ± 0.03. Wilcoxon rank-sum test, p < 0.001. (**c**) Linear classifiers trained on neural responses during initiation epoch successfully decode location from neural responses during the goal epochs. Cross-validated, chance is 25%. pPCx: n = 33 sessions, 8-53 cells/session; CA1: n = 15 sessions, 3-14 cells/session. Each data point corresponds to a single session. Size of data points corresponds to the number of simultaneously recorded neurons. (**d**) Similarity matrix and summary of correlation coefficients for spatial representation between initiation and ITI epochs. r_ppc_same = 0.59 ± 0.03, r_ppc_diff = 0.31 ± 0.03. Wilcoxon rank-sum test, p < 0.001. **(e)** Linear classifiers trained on responses during initiation epoch successfully decode location from neural activity during ITIs. (**f**) Linear classifiers trained on neural activity in correct trials successfully decode location from neural activity during error trials (1 s time window around initiation and goal poke-in were concatenated). (**g**, top) Schematic of maze dimensions. Total footprint of the maze is 1 m x 1 m. Distance between center of port and end of the corridor is 40 cm. (**g**, bottom) Occupancy histogram of relationship between port entry versus distance from port. Grey scale range: 0 - 1 normalized occupancy. (**h**) Activity of neurons for their ‘best corridor’ as a function of distance to port location sorted by peak firing location during a 2.0 s time window prior to initiation port poke-in (before odor onset). Best corridor was defined as the corridor on which neurons had their peak firing. Activity was normalized to the mean firing rate along the corridor. Neurons with firing rates that a peak firing rate ≥ 2x mean rate anywhere on the maze corridor were included (pPCx n = 468 cells; CA1 n = 71 cells). (**i**) Linear classifier decode simultaneously recorded neural population activity along the length of the maze corridor. Cross-validated, chance is 25%. pPCx: n = 33 sessions, 8-53 cells/session; CA1: n = 15 sessions, 3-14 cells/session.

How do spatial representations in pPCx differ from those in the hippocampus? Consistent with the prominent role for hippocampus in spatial navigation, simultaneously recorded CA1 populations in our task more distinctly represented locations as compared to pPCx (**Fig. 2f-k**, 46 CA1 vs. 250 pPCx neuron required to reach ∼ 90% odor decoding accuracy. **Extended Data Fig. 5**). We examined the distribution of peak firing rate for individual pPCx and CA1 neurons as rats traveled across the maze, excluding the periods of odor sampling and reward consumption (**Fig. 3g**). Normalized peak firing locations for CA1 neurons were distributed along the length of the maze arm, while pPCx neurons had peak firing locations at or near ports (**Fig. 3h**). Across simultaneously recorded populations, location classification accuracy for CA1 was comparable along the length of the maze corridor. In comparison, pPCx location decoding accuracy dropped as a function of distance to ports (**Fig. 3i**). This is consistent with features of a learned spatial map which preferentially represented port locations.

### Spatially informative piriform neurons are coupled to the hippocampus

Neural activity across the olfactory-limbic pathway displays dynamical coupling to sniffing or internally generated oscillatory rhythms during behavior^39, 40^. Considering that posterior piriform neurons varied along a spectrum from purely spatial to strict olfactory selectivity (**Fig. 2a-d, Extended Data Fig. 4c, d**), we asked whether this could be due to differences in functional connectivity to either bottom up sensory drive vs. top-down hippocampal network activity. Synchrony of spike timing to sniffing is a signature of neurons in the olfactory system and reflects the feedforward sensory drive from the periphery^3, 41, 42^. Similarly, synchrony of spike timing to hippocampal theta rhythms reflects functional connectivity to hippocampal and parahippocampal networks^43, 44^. We measured sniffing during behavior via a thermocouple implanted in the nasal cavity and the hippocampal theta rhythm by local field potential recording in CA1 (CA1-LFP) (**Fig. 4a**), and used coherence analysis to assess functional connectivity^45^ of pPCx neurons. Sniffing frequency characteristically increased as rats poked their noses into odor ports during odor sampling (**Fig. 4a, Extended Data Fig. 10a, b**)^46, 47^. Location decoding accuracy of pPCx neurons (**Fig. 4b**, bottom) was selectively correlated with the strength of coherence to hippocampal theta and not to sniffing (**Fig. 4c**, bottom row). Odor decoding accuracy of pPCx neurons (**Fig. 4b**, top) was robustly correlated with spike-sniff coherence throughout time (**Fig. 4c**, top), reflecting bottom up sensory drive. Interestingly, odor decoding accuracy was also correlated with CA1-LFP coherence around odor sampling, potentially due to the transient synchrony between sniffing and hippocampal theta band around odor sampling (**Extended Data Fig. 10c**). The coupling of location-selective cells to hippocampal theta and odor-selective cells to sniffing was reproduced in an analysis of the difference in coherence for the best odor-decoding vs. the best location-decoding neurons (**Extended Data Fig. 11**). Consistent with functionally distinct subpopulations of pPCx neurons, we also found that noise correlations were higher within odor-and location-selective subpopulations than between (**Extended Data Fig. 9**). We interpret these results to suggest that subpopulations of piriform neurons have differential connectivity to either the olfactory bulb or the hippocampal network^23, 48–50^, as well as stronger within-population coupling. However, given that we observed a spectrum of response properties ranging from purely sensory to strictly spatial, these subpopulations are likely not strictly segregated.

**Figure 4.**
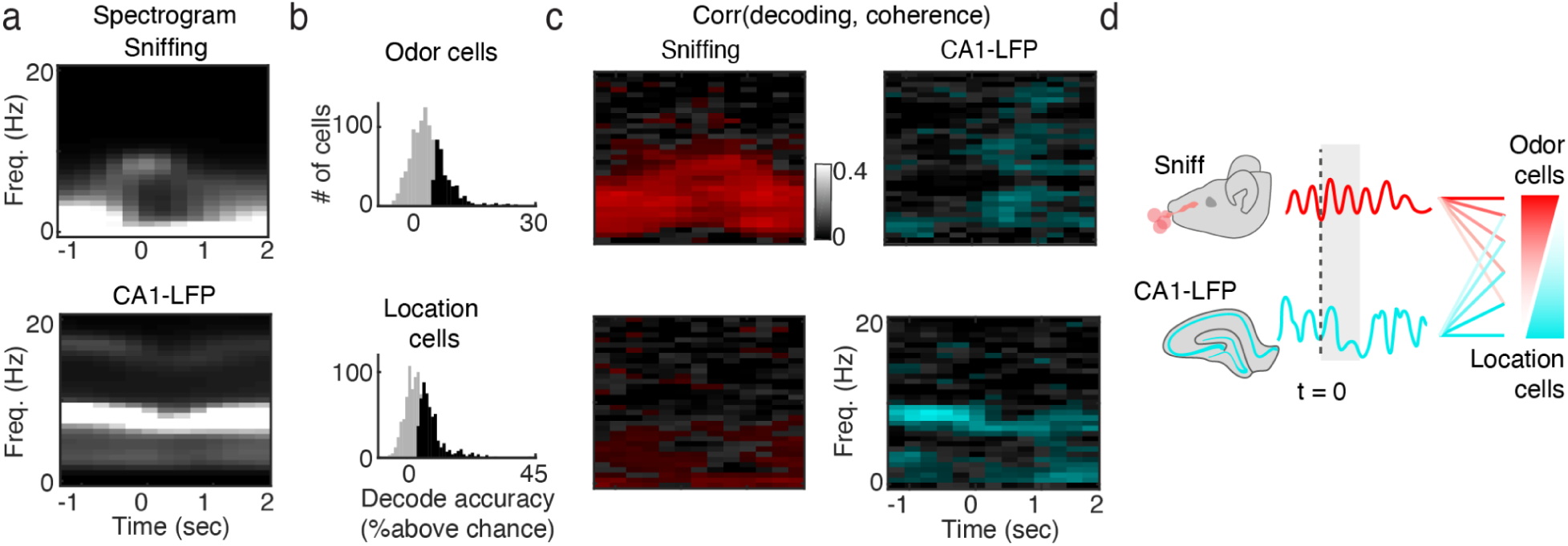
Location neurons are coupled to hippocampal theta rhythms. (**a**) Average spectrogram of sniffing (top) and CA1 LFP (bottom) aligned to odor onset time. Brightness from 0 to 90th percentile of power in each panel. (**b**) Decoding performance of pPCx neurons for odor (top) and location (bottom). Significance was defined as accuracy greater than the 95th percentile of classifiers trained on shuffled labels. (**c, top**) Correlation between odor decoding performance and coherence of spikes to sniffing (left) and CA1-LFP (right). Colored pixels indicate frequency-time bins in which decoding performance was significantly correlated with sniffing (red) and CA1-LFP (cyan). The coherograms of cells identified as significantly odor (location) coding (Fig 4b) were taken, and for each (time, frequency) value of the coherogram, the across-cell correlation between coherence and decoder was calculated. To identify (time, frequency) pairs with a significant association between decoding and coherence, the correlation was repeated 1,000x using shuffled decoder values, and (time, frequency) pairs whose value exceeded the 95th percentile were colored red (for sniff) and cyan (for CA1-LFP). (**c, bottom**) Similar analysis as (c, top) but for location decoding accuracy. (**d**) Schematic of preferential coupling between odor and location cells to sniffing and CA1-LFP, respectively.

## Discussion

In this study, we examined the function of the rodent piriform cortex while rats used odor cues for spatial choice and found that populations of posterior piriform neurons fired in a spatially restricted manner in the environment. Spatial representations in the pPCx resemble hippocampal cognitive maps, as they were stable independently of immediate sensory drive, reward prediction, or task engagement.

As compared to aPCx^3, 30, 32^, a relatively large number of pPCx neurons were required to decode odor identity. The relatively slow rise time and long integration time window of pPCx odor decoding also contrasts sharply with the fast onset and tightly respiratory locked spike times of aPCx neurons^3^. We interpret these findings as evidence for a functional distinction between these two regions of piriform^20, 21, 23, 51^, where pPCx is comparatively less dedicated to odor coding. In contrast to the modest decoding performance for odors, we found that pPCx robustly encoded port locations in our task, and location-informative neurons were strongly coupled to CA1-LFP at theta-band frequency. Learning and shifting behavioral demands shape computations performed by brain regions. In our task, the odor discrimination aspect was not very demanding: animals were presented with a fixed set of four distinct monomolecular odorants. We speculate that given a more dynamic and complex olfactory environment pPCx could be more engaged in odor coding.

We note that while projections from hippocampus to piriform cortex are weak, projections from medial and lateral entorhinal cortices^49, 50, 52, 53^ provide a direct pathway for top-down influence on the piriform. The coupling of pPCx neurons to sniffing rhythms and CA1-LFP reflects the convergence of bottom-up sensory versus top-down cognitive drive to this region. This convergence likely results in the functional heterogeneity of pPCx neurons: neurons range from purely odor-selective and sniffing-coupled neurons to spatially-selective and theta-coupled. Neurons in the middle of the spectrum with dual odor- and spatial-selectivity, presumably receiving converging bottom-up and top-down information, are well-positioned to connect transient sensory cues to spatial goals during navigation. Future studies on the distinct functions of aPCx and pPCx, as well as subregions of olfactory cortex outside of piriform cortex such as the anterior olfactory nucleus or cortical amydala^23, 32, 33, 51, 54, 55^ will undoubtedly contribute to a more comprehensive understanding of integration of olfactory and spatial information, and the distinct roles of neural pathways engaged during odor-guided spatial behaviors.

How could piriform spatial maps be generated? Sensory projections from the olfactory bulb to piriform cortex are thought to be largely unstructured^51, 56–59^, and piriform recurrent connections are plastic, broadly distributed and lacking in topographical organization^12–16, 60^. These circuit features are characteristic of learning systems^13, 17^, and active local mechanisms within piriform likely work in concert with top down projections^23, 48–50^ to give rise to the spatial representations we observed. Indeed, while firing fields of CA1 place cells tile the length of the maze corridor, spatially-informative pPCx neurons cluster around port locations. This is consistent with an underlying associative mechanism linking odor cues to spatial locations in pPCx. Given that, in our task, water rewards were delivered at the same location as odors, it is not clear whether clustering of spatial representations at ports reflects purely odor-place associations or whether rewards also contribute to map formation.

Cognitive maps are internal models used to represent predictable structures in the world, and reflect an animal’s prior knowledge^34, 38, 61, 62^. Animals can produce adaptive and flexible behavior by combining incoming sensory evidence with internally-held cognitive maps of the environment. What could be the functional distinction between hippocampal versus pPCx representations? Neurons in the hippocampal formation respond to sensory cues, including odors^63–68^, and sensory cues can also induce global firing rate changes in hippocampal representations referred to as “remapping”^69^. We speculate that while the hippocampal formation may provide a complete and continuous cognitive map of the spatial environment that is modulated by sensory input^34, 38, 70^, pPCx could serve to represent select locations in the environment relevant for olfactory-driven behaviors.

A longstanding question in neuroscience is the neural mechanism by which sensory evidence interacts with these abstract cognitive variables to produce behavior. Such abstract cognitive variables are thought to be generated by and held in higher order brain regions, such as prefrontal cortex or the hippocampal formation^61, 62^. The representation encoding of odors and space in piriform illustrates that abstract cognitive variables can be prominently encoded by sensory regions outside of the canonical circuits for spatial cognition. It is likely that the posterior piriform cortex may serve as an odor-driven learning system^71^ more akin to the hippocampus than to canonical primary sensory cortices, possessing computational and circuit properties of both sensory and learning systems. We posit that our work provides a new avenue of inquiry on the fundamental question of how sensory information is combined with cognitive maps in the brain to guide flexible behaviors.

## Methods Summary

All experiments and procedures were approved by the Champalimaud Foundation Bioethics Committee and the Portuguese National Authority for Animal Health, Direcção-Geral de Alimentação Veterinária (DGAV). See Full Methods.

## Acknowledgements

We are grateful to J. J. Paton, B. A. Atallah, L. L. Glickfeld, and J. B. Hales for comments on the manuscript; E. Lottem, M. Murakami, M. G. Bergomi, C. Linster, A. Fleischmann, K. M. Franks, S. R. Datta, C. E. Schoonover, A. J. P. Fink, and R. Axel for helpful discussions. A. S. Cruz and A. C. Rato for assistance with animal training; L. M. Frank and members of the Frank laboratory for experimental advice and assistance; Champalimaud Research Hardware Platform for custom components used in the behavioral task and recordings; Champalimaud Foundation, European Research Council(Advanced Investigator Grant 671251), Human Frontier Science Program(LT000402/2012), Fundação para a Ciência e a Tecnologia (FCT-PTDC/MED-NEU/28509/2017, C.P.), and Helen Hay Whitney Foundation for financial support.

## Author Contribution

The project was originally conceptualized by C.P. and Z.M. and further developed in collaboration with N.B. The behavioral paradigm was developed and designed by C.P., N.B, and Z.M. Task-related hardware was developed and constructed by N.B. and C.P. Task-related software was developed and implemented by N.B. Animal training, behavior data collection, and behavioral data analysis was performed by C.P., and N.B. Neural recordings were performed by C.P. Neural data analysis was performed by C.P. and G.A. The manuscript was written by C.P., G.A., Z.M. and edited and reviewed by all authors.

## Competing interests

The authors declare no competing interests.

## Extended Data Figures 1 - 11

**Extended Data Figure 1.**
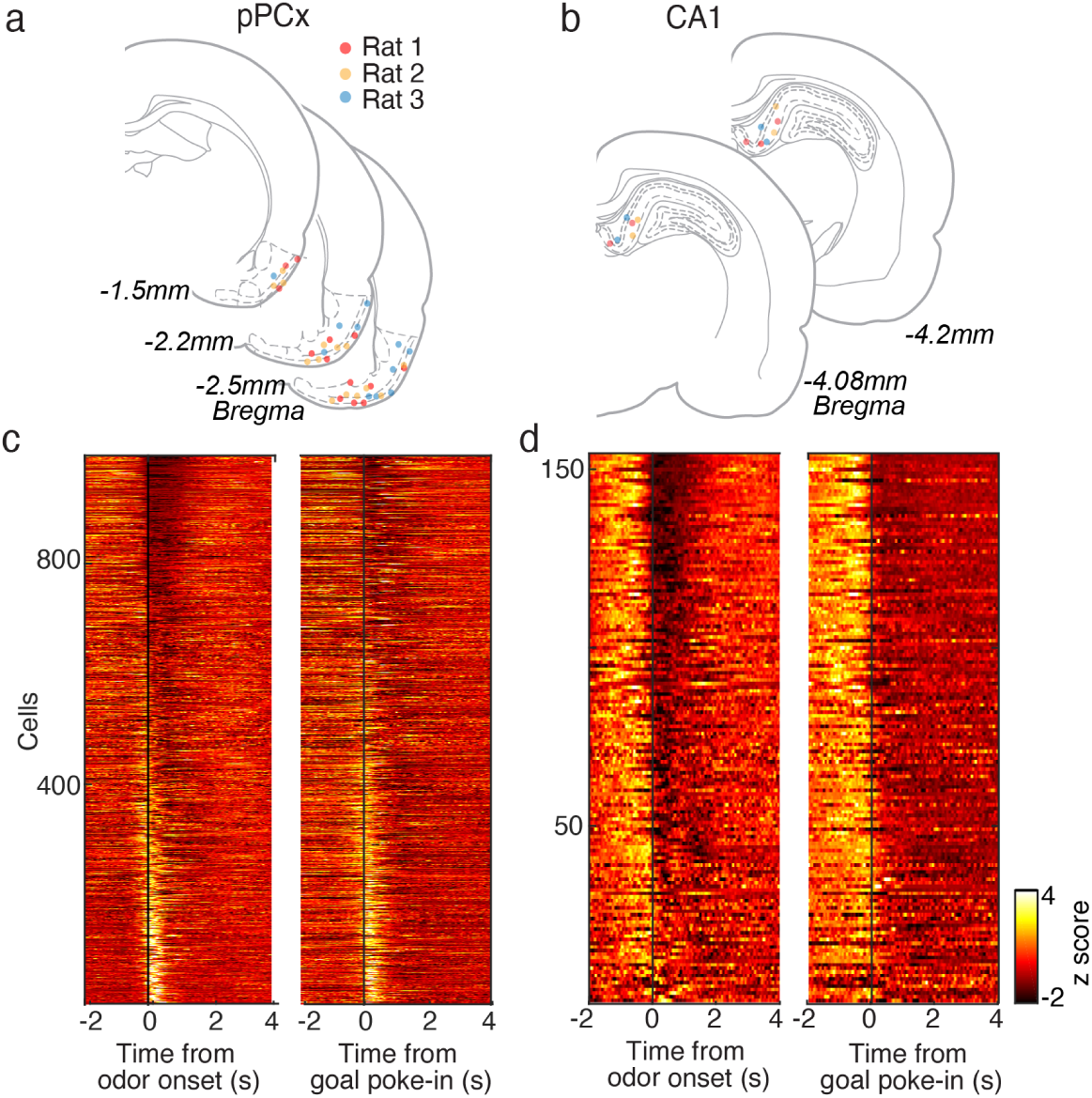
Tetrode lesion sites and population summary of pPCx and CA1 neurons. (**a**) Tetrode lesion sites for 3 recorded rats in pPCx. See full methods for targeting and verification of tetrode recording sites. Due to the wide range of tetrode lesion sites (from -1.5 mm to -2.5 mm bregma), lesion sites from multiple histological sections were summarized onto 3 representative atlas sections. (**b**) Summary PETH for all pPCx (n = 995) neurons recorded. Spike timing was aligned to odor onset (left) and goal poke-in (right). Neurons were sorted based on their activity during 1 s time window after odor onset in both left and right panels. For each neuron, the mean z-scored rate during a 2 s time window prior to alignment time point was subtracted from the entire PETH. (**c**) Tetrode lesion sites for 3 recorded rats in dorsal CA1. See full methods for targeting and verification of tetrode recording sites. (**d**) Summary PETH for all CA1 (n = 154) neurons recorded.

**Extended Data Figure 2.**
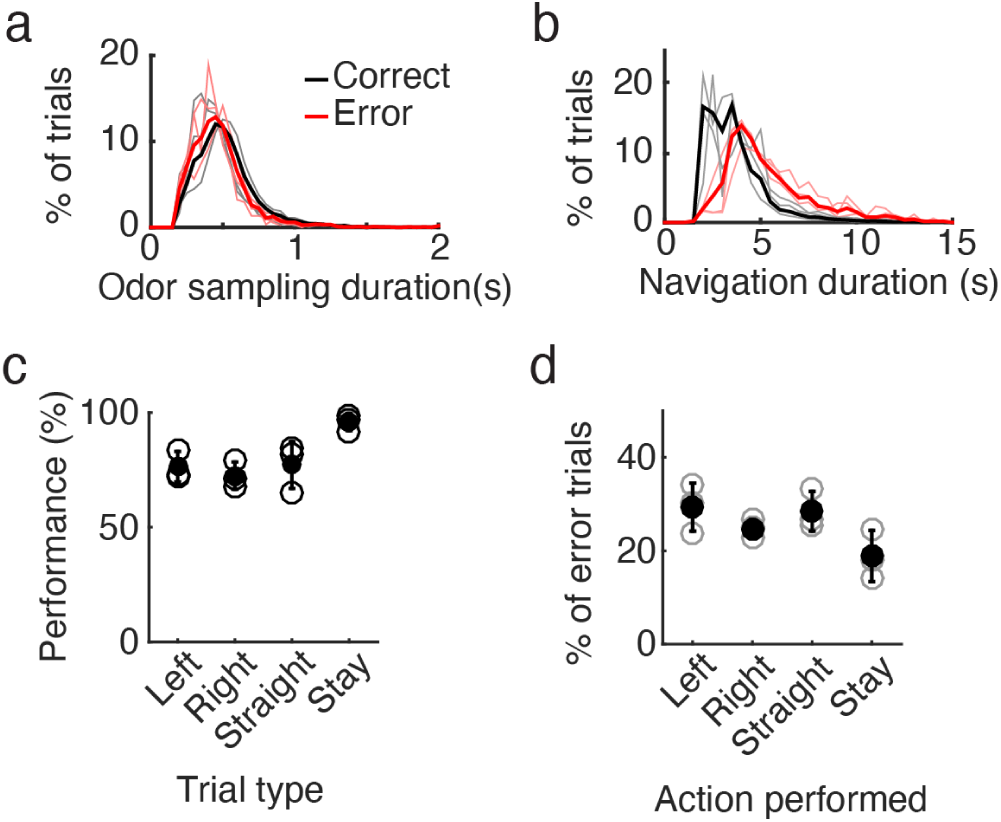
Correct and error trial behavior in odor-cued spatial choice task. (**a**) Odor sampling duration in 3 recorded rats. A minimum of 150 ms of odor sampling was enforced. Black: correct trials, 0.519 ± 0.02 s. Red: error trials, 0. 448 ± 0.04 s. (**b**) Navigation duration in 3 recorded rats, as defined by time between initiation (odor) port poke-out and goal port poke-in. Goal ports were only active (reward available) after a 1.5 s delay after rats poke out of initiation ports. Black: correct trials, 3.67 ± 0.15 s. Red: error trials, 5.35 ± 0.8 s. (**c**) Rats performed better for trials in which goal location indicated by the odor cue was congruent with odor sampling location (stay trials). n = 3 rats, ANOVA, p = 0.01, Mean ± S.D. (**d**) Actions performed for error trials. ANOVA, p = 0.07. Mean ± S.D..

**Extended Data Figure 3.**
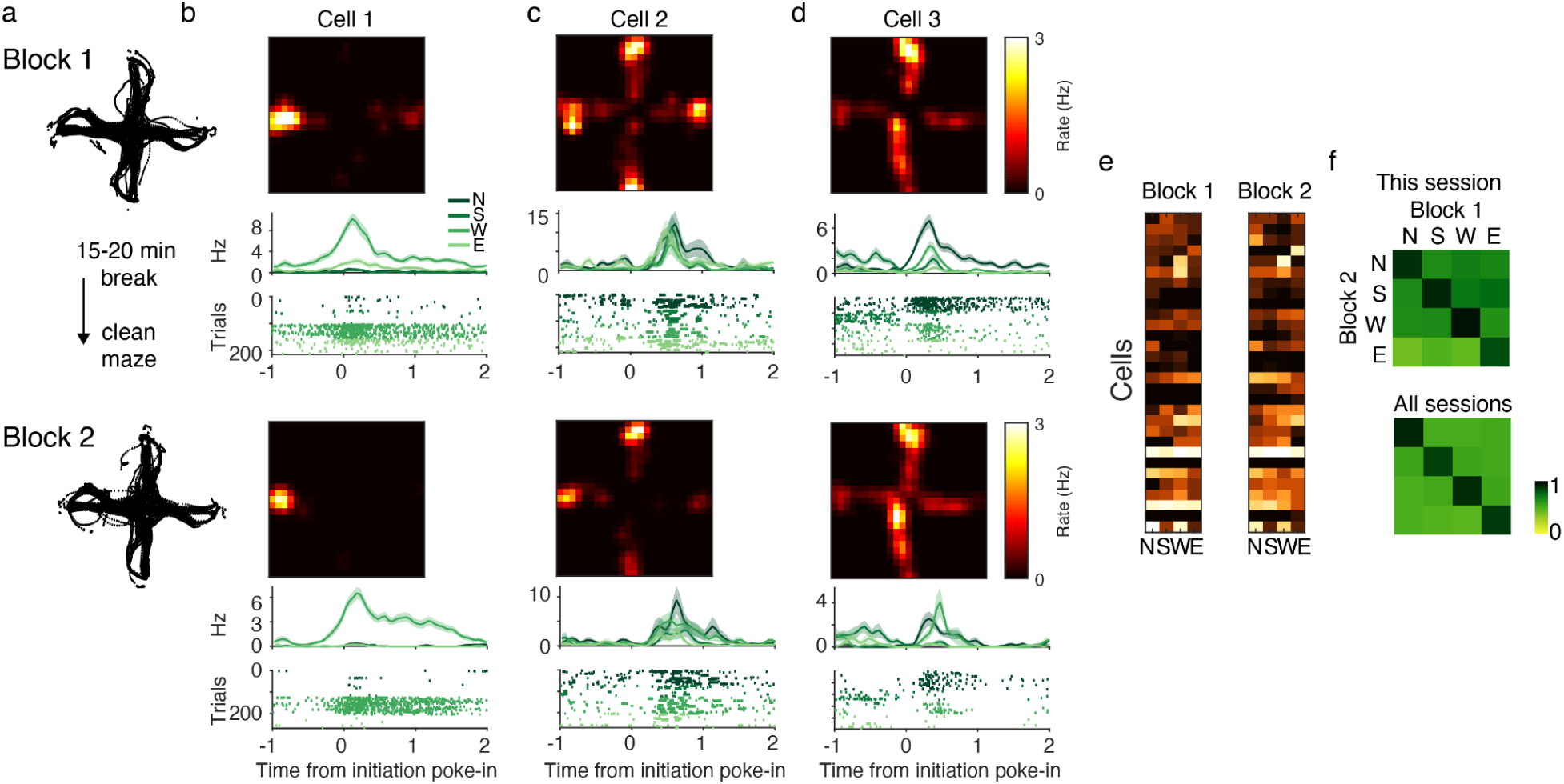
pPCx location selectivity remains stable across recording blocks. (**a**) Rat position on the maze during two blocks within the same recording session. (**b-d**) Occupancy normalized firing rate heat map, PETH, and raster plots associated with example cells 1-3 in Fig 2a-c. PETH and raster plots were aligned to initiation port poke-in and sorted by location. (**e**). Z-scored mean firing rate of neurons for different port locations across blocks. (**f, top**) Similarity matrix obtained by computing the pairwise Pearson’s correlation coefficients between locations population response vectors for blocks 1 and 2 of example session shown in (f). Coefficients along the diagonal band are significantly higher than off-diagonal (r_diag = 0.80 ± 0.05, r_offdiag = 0.54 ± 0.09, Wilcoxon rank-sum test, p < 0.01), representing stability of location representation across blocks. (**f, bottom**). Same analysis as in the left panel but for all pPCx neurons recorded (r_diag = 0.81 ± 0.02, r_offdiag = 0.47 ± 0.02, Wilcoxon rank sum test, p < 0.001, n = 44 sessions, 995 neurons).

**Extended Data Figure 4.**
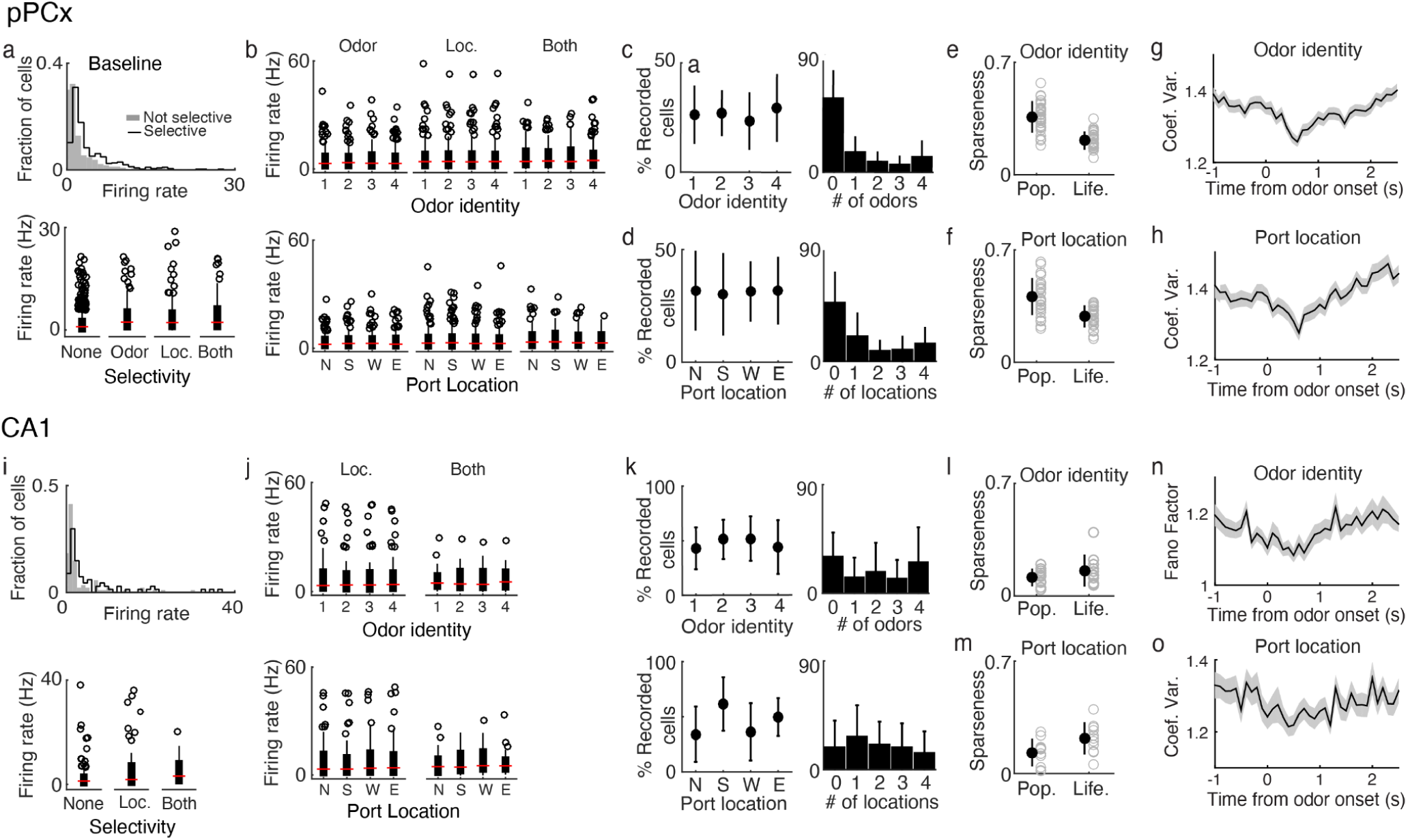
Summary of pPCx and CA1 response properties for odors and port locations in task. (**a**) Firing rate for individual pPCx neurons throughout recording sessions (Non-selective neurons: median = 0.94, range = 0.03 - 21.48 Hz, n = 531 neurons; selective neurons (for either odor or location): median = 1.95 Hz, range = 0.04 - 27.49 Hz, n = 531 neurons; Wilcox rank-sum test p < 0.01). (**a**, bottom) Firing rate of neurons grouped by selectivity properties. Odor-selective only neurons, n = 120 neurons, median = 1.93, range = 0.05 - 17.84 Hz. Location-selective only neurons, n= 238, median = 2.06, range = 0.04 - 27.49 Hz. Odor and location selective neurons (joint): n = 106, median = 1.88, range = 0.14 - 17.41 Hz. (**b**, **top)** Firing rates for individual neurons across 4 odor identities during 0 - 1.0 s post odor onset. Neurons are grouped by selectivity properties. Red tick: median; edges of the bar indicate the 25th and 75th percentiles; circles: outliers. Odor: neurons selective to odor identity. Loc: neurons selective to port locations. Both: neurons selective to both odors and port locations. **(b, bottom)** Same as top but across odor sampling port locations. (**c**, **left)** Fraction of recorded pPCx neurons in a session that was activated by different odors. (**c**, right) Histogram of number of odors that activate pPCx cells in a session. Activation was measured by comparing mean firing rate for 1 s after odor onset time compared to a 2 s baseline immediately preceding odor onset (n = 33 sessions). Wilcoxon rank-sum test, p < 0.05, corrected for multiple comparisons (Full methods). (**d**) Similar to (d) but for port locations. (**e**) Sparseness across odor identity for simultaneously recorded populations. Population sparseness = 0.36 ± 0.09; lifetime sparseness = 0.21 ± 0.05, Mean ± S.D. (n = 33 sessions). (**f**) Similar to (d) but for port locations. Population sparseness = 0.40 ± 0.12; lifetime sparseness = 0.29 ± 0.06. (**g**) Coefficient of variation for odors aligned to odor onset. Trial-to-trial coefficient of variation for each neuron was calculated for different odor trials and averaged. (**h**) Similar to (g) but calculated for different port locations. (**i-o**) Same analysis as (**a-h**) for CA1 population.

**Extended Data Fig 5.**
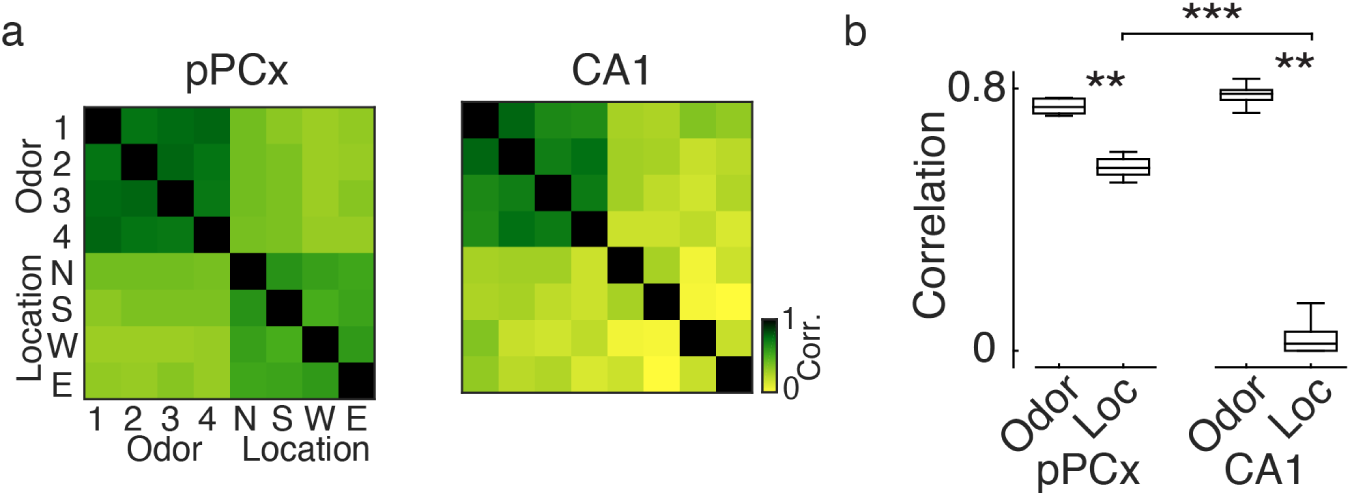
(**a**) Full similarity matrix of correlation coefficients of all odor and location population vector pairs for all recorded pPCx (n = 995) and CA1 (n = 154) neurons. Off-diagonal correlation coefficients on the lower right quadrants of each matrix show that CA1 location representations are more dissimilar from each other than pPCx location representations. Off-diagonal correlation coefficients on the top right and lower left quadrants show that there were no systematic relationship population responses for individual odors and locations. (**b**) Population correlation coefficients for odor and locations are shown in similarity matrices. Odor: top left quadrant; location: bottom right quadrant, excluding autocorrelation coefficients (diagonal band). Wilcoxon rank-sum test, ** p < 0.01, *** p < 0.001.

**Extended Data Figure 6.**
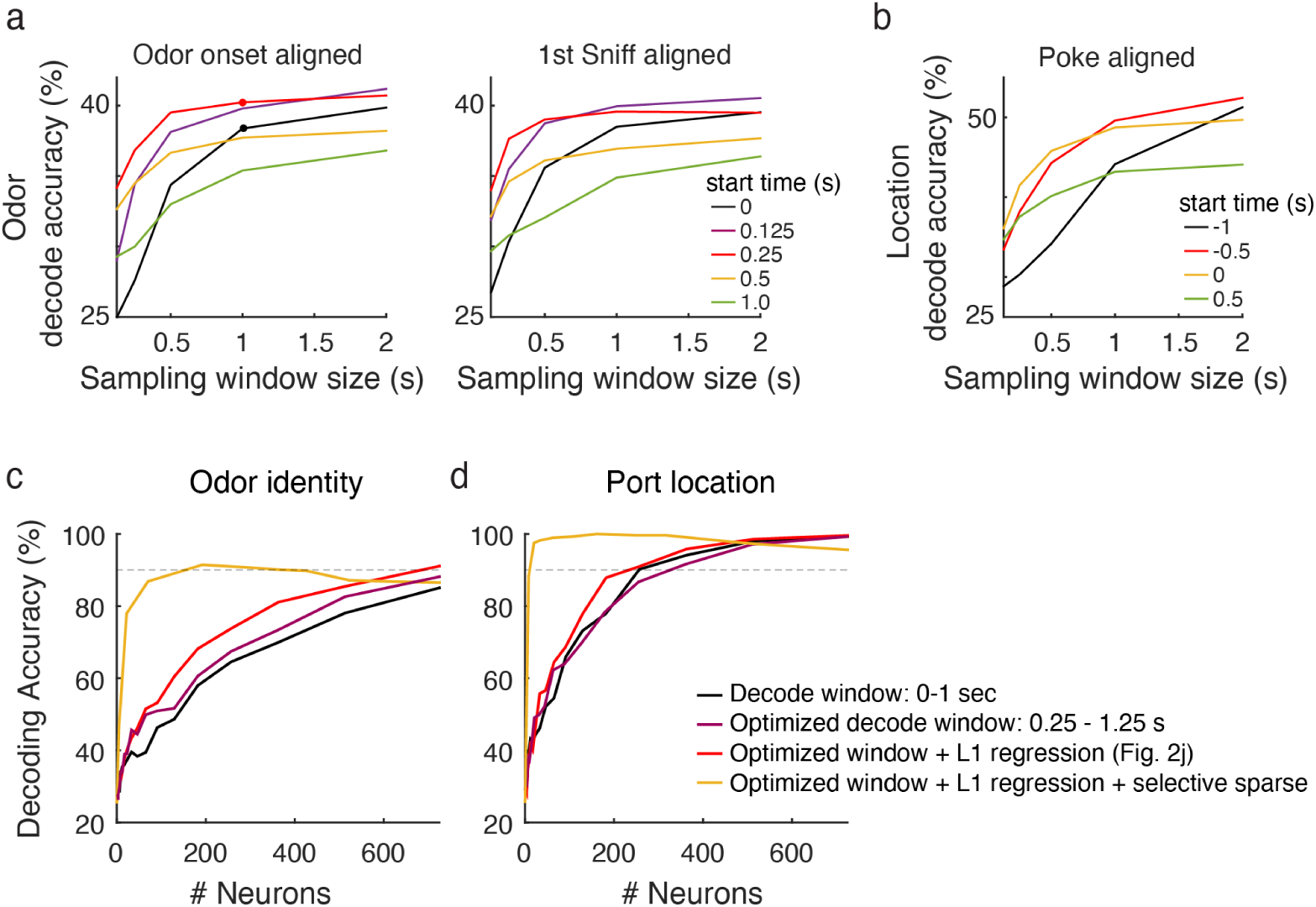
Optimization of pseudo-population decoding parameters. (a) Odor decoding accuracy for a simultaneously recorded pPCx population (example session from Figure 2) using a wide range of time windows aligned to odor onset time (left), or first respiration after odor onset (right). The black and red dots indicate time windows used for black and red lines in (c), respectively. (**b**) Location decoding accuracy aligned to initiation port poke-in time across a wide range of time windows. (**c**) Pseudo-population decoding of odor identity with different time windows and regularization. Black and purple lines use L2 regularization, while red and yellow use L1 (sparse) regularization. By increasing the sparsity of the L1-decoder (scanning the ‘cost’ parameter over the range 2^[-7:8]) and plotting decoding accuracy as a function of the number of contributing neurons (# of neurons with non-zero weights in the decoder), we can minimize the contribution of uninformative neurons. Here, the x-axis indicates the number of neurons being used by the decoder (i.e. neurons with non-zero kernel weights), which was controlled by changing the L1 penalty. When this penalty is large, the decoder selectively uses only the most informative neurons, leading to a much steeper rise than seen for the L2 regularization pseudo-population curves, which sample neurons randomly. Using this sparse L1-decoding approach, it is clear that odor identity can accurately be decoded from a small population of pPCx neurons (90% decoding accuracy for ∼150 neurons). Dotted line indicates 90% accuracy. Red line is the same as dashed line in Fig. 2g. **(d)** Same analysis as in (c) but for port locations. Red line is the same as dashed line in Fig. 2k. Note that while it conveys that relatively few neurons are needed to encode odor information, the steepness of the yellow curve is sensitive to the number of recorded neurons (since a larger pool is more likely to contain an informative neuron).

**Extended Data Figure 7.**
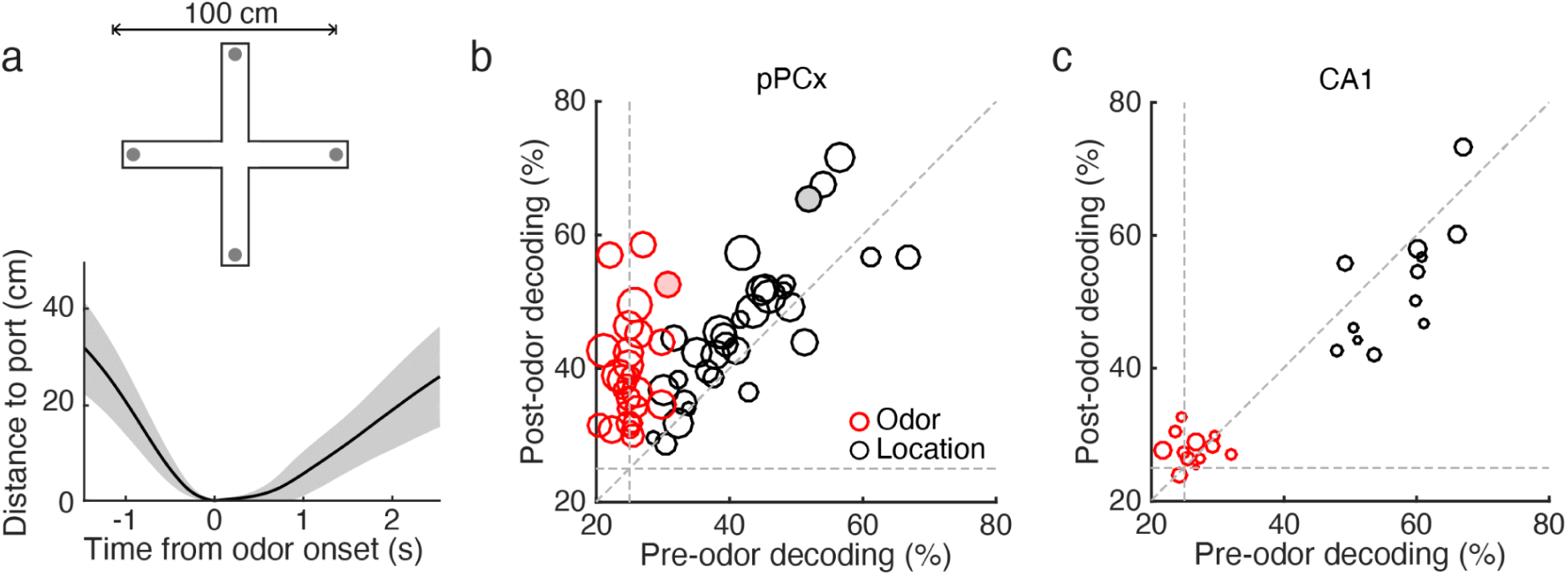
Location decoding accuracy is independent of olfactory drive. (**a**) Distance from port locations aligned to odor onset. (**b**) Population decoding accuracy for odor and port locations across time for simultaneously recorded pPCx (n = 33 sessions; 8-53 cells/session) . Pre-odor decoding used population activity from 1.5 s before odor onset; post-odor decoding used population activity 1.5 s after odor onset. Cross-validated, chance is 25%. The filled-in data points indicate the example session shown in Figure 2. (c) Same analysis as in (b) for simultaneously recorded CA1 population (n=12 sessions, 3-14 neurons/session).

**Extended Data Figure 8.**
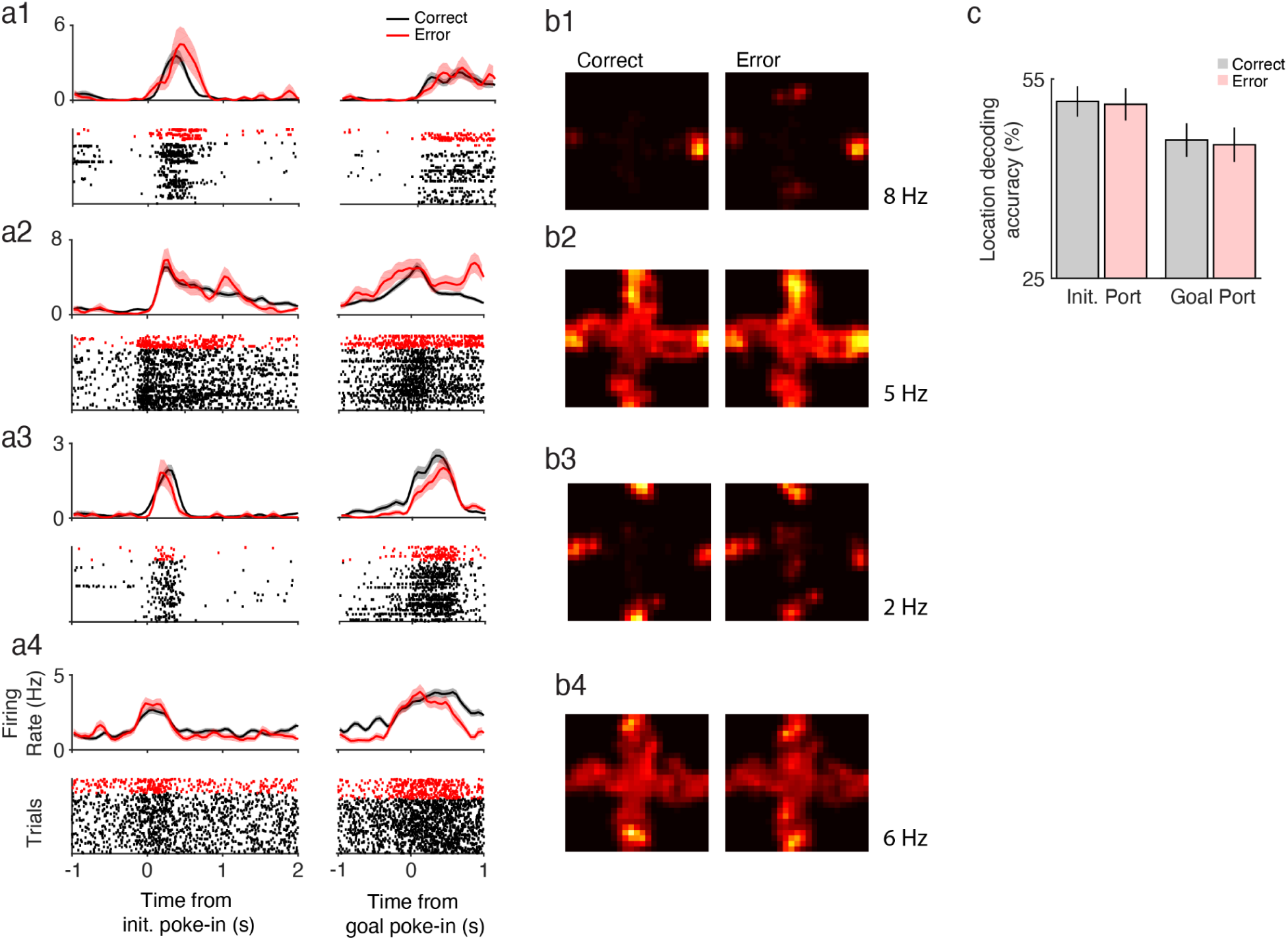
Example cells and location decoding for correct and error trials. (**a1-4**) PETH and rastors for 4 example neurons aligned to initiation port poke-in (left) and goal poke-in time (right). Black: correct trials; red: error trials. (**b1-4**) Firing rate heat maps for example cells. Heat maps were normalized to occupancy and generated by concatenating all trials from -1 to 2 s window around initiation and goal port entry. Peak firing rates noted to the right of heat maps. (**c**) Location decoding accuracy of a classifier trained on neural activity -0.5 to 0.5 s around initiation port poke-in for correct trials. The classifier was tested on neural activity -0.5 to 0.5 s around initiation and goal port poke-ins for correct (black) and error (red) trials (cross-validated, mean ± S.E.M., n = 33 sessions; 30 ± 13 neurons/session).

**Extended Data Figure 9.**
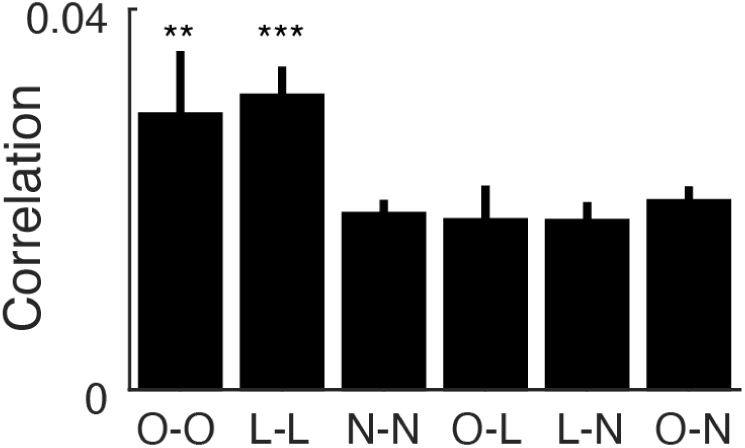
Noise correlations between pPCx neurons are consistent with distinct functional subgroups. Noise correlations between 3 groups of neurons (6 group pairings): odor-selective (O), location-selective (L), non-selective (N) neurons. Overall noise correlation between groups increased with longer time windows. O-O and L-L correlations are higher than for N-N. Mean ± S.E.M., ** p < 0.01, *** p < 0.001.

**Extended Data Fig 10.**
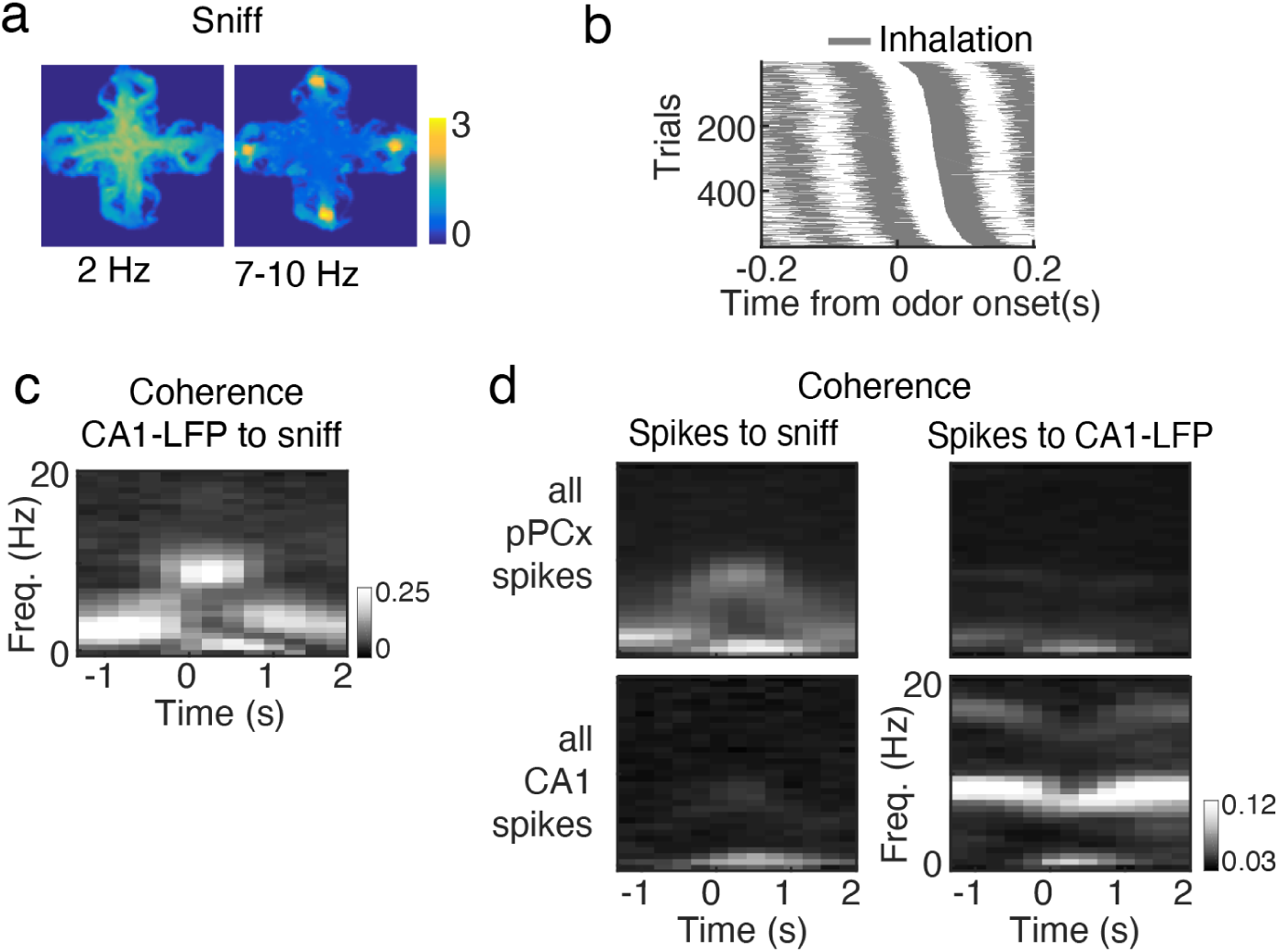
Sniffing and hippocampal theta-band activity in task. (**a**) Sniffing behavior on maze in one example session. Heat map of basal sniffing behavior (2 Hz power) and high frequency active sniffing behavior (7-10Hz power) on maze. Colorbar is power. (**b**) Sniffing behavior for one example session. Sniff phase was time-locked to odor port poke-in. Gray color is inhalation, white is exhalation. (**c**) Coherence between CA1-LFP and sniffing. Time is aligned to odor onset. Colorbar is coherence. (**d, top**) Average coherence of spike times for all pPCx neurons to sniff (left) and CA1-LFP (right). Time is aligned to odor onset time. (**d, bottom**) Average coherence of spike times for all recorded CA1 neurons to sniffing (left) and CA1-LFP (right).

**Extended Data Figure 11.**
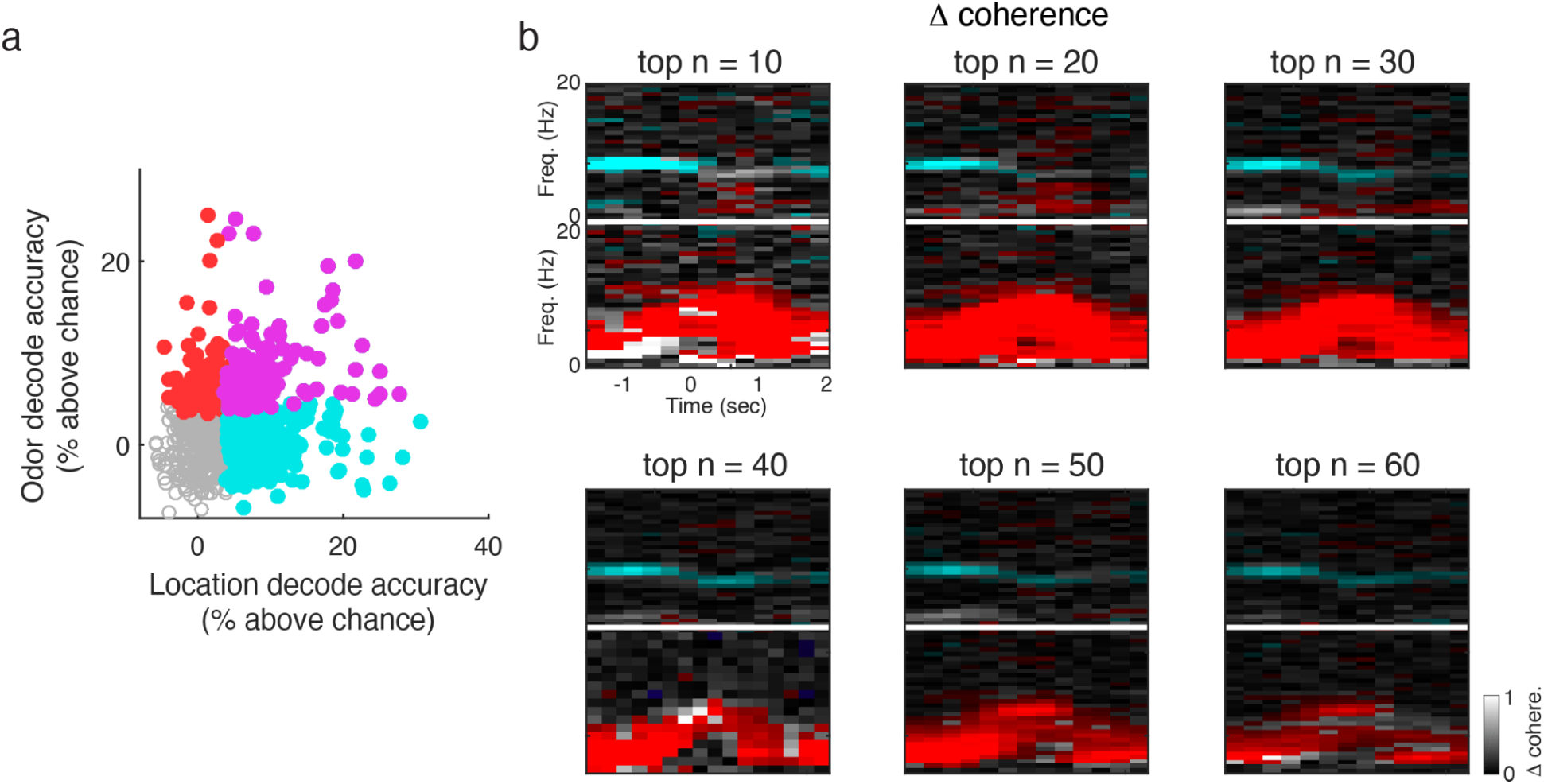
Preferential coupling of odor cells to sniffing and location cells to hippocampal theta. (a) Odor and location decoding accuracy of individual neurons. Neurons decoded odor only (red), location only (cyan), both (magenta), or neither (open circles). Decoding significance was defined as accuracy greater than the 95th percentile of classifiers trained on shuffled labels. **(b)** Mean difference in coherence between the ‘top n’ best odor decoding neuron vs ‘top n’ best location decoding neuron. Top panels of each ‘top n’ analysis show differences in spike-sniff coherence, while bottom panels show differences in spikes-to-CA1-LFP coherence. Red pixels indicate frequency-time bins in which spikes from odor decoding cells are significantly more coherent than spikes from location decoding neurons, while blue pixels indicate the bins in which location cells are more significantly coherent than odor cells. Gray scale is coherence. (see Full Methods).

## Full Methods

### Animal Subjects

A total of 6 male adult Long-Evans hooded rats (4–6 months and weighing 450–550 g) were used in experiments. All 6 rats were used for behavioral analysis (**Fig. 1**), 3 rats were implanted for neural recordings (**Fig. 2 to 4 and all Extended Data Figs**).

### Odor-cued spatial navigation task

Rats were trained and tested on an odor-cued four-alternative spatial choice task where water was used as a reward based on tasks previously described^1^. Here, rats were trained on an elevated plus maze with open arms and a 25mm ledge. At the end of each arm there was one nose port which could deliver a light cue, odors, or water reward (modified from Island Motion)^2^. Each odor was associated with one possible reward location (North, South, West or East) (**Fig 2a-c**), each trial followed the structure of initiation/odor delivery, navigation, and reward. A trial begins with a light cue indicating the initiation port for that trial. Rats initiated a trial by nose-poking at the port indicated by a light cue. This triggered the delivery of an odor with a (uniform) random delay of 0.1–0.25 s. Initiation port location was pseudo-randomized within 10 trial blocks and balanced across all port locations. Rats were trained to stay in the odor sampling port for a minimum of 0.15s, after which they were free to leave (**Extended Data Fig. 2a**). Odor delivery was terminated as soon as the rat exited the odor port. Following poke-out of the initiation port, a minimum 1.5 second time delay for choice/navigation is enforced to discourage rats from their preference to stay at their current location to collect reward (**Extended Data Fig. 2b**). After this delay period, water reward is available at the correct port location. A poke in the correct location would yield a tone (80ms 3KHz) and a 30µl water reward while a poke in an incorrect port would yield an error tone (80ms white noise burst). A 4 to 6 seconds (uniform distribution) inter-trial interval (ITI) period started after the delivery of the reward. Reward was available for correct choices for up to 10 s after the rat left the odor sampling port. Water was delivered with a random delay from entry into the goal port drawn from a uniform distribution of 0.5-1.0 s. The task was designed and implemented using a RTLFSM and B-control as described previously ^3^. A Point Gray Flea3 1.3MP camera was used to track the rat’s behavior. The Bonsai framework ^4^ was used to interface with the cameras.

### Odor stimuli

Odor delivery was controlled by a custom made olfactometer^3^ calibrated by a photoionization detector (miniPID, Aurora Scientific). We used relatively low concentration of odorants (liquid dilution factor: 1:100 in mineral oil and further diluting 100 ml/min odorized air in a total of 1000 ml/min clean air stream). All odor stimuli (1-Hexanol, Caproic Acid, R-Limonene, and Amyl Acetate) were randomly interleaved during a session.

### Training

Rats had free access to food, but water was restricted to the training sessions and 5-10 additional minutes of free access per day. Rats were trained in 45min sessions, twice a day, 5-7 days/week. Each training session was spaced 6-8 hours apart for motivation. To prevent over-training, in 20% of trials (pseudo-randomly interleaved), the correct choice is indicated by a light cue at the correct goal port (answer trials). In the remaining 80% of trials, all 4 possible goal ports are lit (question trials). A threshold of 75% correct for question trials (chance is 25%) is taken as performance criterion. All behavioral and neural data analysis were restricted to question trials only.

### Microdrive implant

After reaching an asymptotic performance in behavioral training, each rat was implanted with a custom-designed and 3D printed multielectrode drive (based on microdrives designed in the laboratory of Dr. Loren Frank, U.C.S.F) (PolyJetHD Blue, Stratasys Ltd.) with 24 independently moveable tetrodes (based on design from the laboratory of Dr. Loren M. Frank., UCSF). Tetrodes (Ni-Cr, California Fine Wire Company) were arranged into a 3 x 8 array within a cannula and gold plated to reach an impedance of 250 kΩ at 1 kHz. Implanted recording drives had two cannulas that targeted both pPCx (19 tetrodes, rectangular cannula angled at 19 degrees coronal angle away from the midline), and dorsal CA1 (5 tetrodes, vertical cannula). Cannula for pPCx was positioned such that tetrodes spanned anterior to posterior range of piriform cortex. Cannulas were centered at the following coordinates: right pPC -2.0 mm AP and 3.0 mm ML, right dorsal hippocampal CA1 −4.2 mm AP and 2.0 mm ML. LEDs were attached to the microdrive and used for video tracking of rat position.

### Neural recordings

After 10 days of post-operative care and recovery, rats were water restricted and trained in the same manner as the pre-surgery period. Tetrodes were adjusted every 2 days post-surgery to reach the target coordinate, guided by depth, LFP, and spiking patterns. Before starting recordings, animals were retrained to reach similar accuracy levels as those achieved before surgery (>75% correct for question trials). Recording sessions were split into two 45min behavioral blocks with a 15-20min rest period in between blocks to allow for rats to rest and consume food in a separate box. During this time, surfaces of the maze and port were cleaned thoroughly with enzymatic cleaner (Henry Schein Medical) and disinfectant (VirkonTM-S, and 70% ethanol). Electrical signals were amplified and recorded using the Cerebus data acquisition system (Blackrock Microsystems, Utah). Thresholded events were recorded at 20kHz (for spike sorting), continuous signal was recorded at 2kHz (for LFP). Due to the large distance between tetrode entry point and target piriform cell body later, in order to target piriform primary cell body layer, a single tetrode in the most anterior position was advanced ahead of the rest (‘scout’ tetrode), past the entire principal cell body layer (as identified by increases in threshold-crossing events), to reach the ventral lateral inner surface of the cranium (as identified by a signature pop of voltage saturation followed by complete lack of threshold-crossing events). The remaining tetrodes were then advanced to the cell body layer depth as identified by the ‘scout’ tetrode, while taking into account the medial-lateral inclination of cell body layer depth. Tetrode depths were adjusted at the end of each session while monitoring spike waveform and firing properties in order to sample an independent population of neurons across sessions. The locations of tetrode tips during each recording session were estimated based on their depth and histological examination based on electrolytic lesions and the visible tetrode tracks. Rats performed 1 session per day, and a total of 44 recording sessions were obtained from 3 rats.

### Recording and analysis of respiration pattern

To monitor sniffing, during drive implantation, a temperature sensor (custom T-type probe, 44ga., Physitemp Instruments) was implanted in one nostril, and respiration patterns were monitored as a temperature change in the nasal cavity as described previously ^3^. Signals from nasal thermocouple were amplified, filtered between 0.1 and 475 Hz and digitized at 2,000 Hz. For analysis, voltage signals were further low-pass filtered (< 50 Hz). Onset of inhalations and exhalations were identified as local maxima and minima of the signals semi-automatically using custom software.

### Histology

To verify the ultimate location of the tetrodes, electrolytic lesions were produced after the final recording session (30 μA of cathodal current, 3 s). Rats were deeply anesthetized with pentobarbital and perfused transcardially with 4% paraformaldehyde (wt/vol in PBS). The brain was sectioned at 50 μm and stained with Cresyl violet solution to observe sites of electrolytic lesions (**Extended Data Fig. 1**).

### Data Analysis and Statistics

All data analysis and statistical tests were performed with custom-written software using MATLAB (Mathworks). No statistical methods were used to pre-determine sample sizes.

#### Behavioral bias analysis (Fig. 1g, h)

Delta bias is the change in probability of choosing a particular location in the next trial as a consequence of the outcome of the current trial. We can express this as:

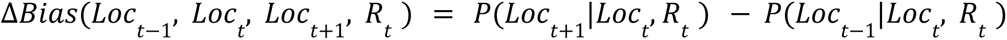

Where **t** is the current trial; location **Loc** ∈ {North, South, West, East} and reward **R** ∈ {0,1} . The Location bias is defined as the delta bias of repeating the same location and results from fixing the Location and conditioning the analysis to rewarded and non rewarded trials:

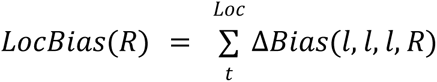

The same analysis was performed for actions by substituting Loc for Action ∈ {Left, Right, Forward, Stay}.

#### Spike Sorting

Single units were isolated offline by manually clustering spike features derived from the waveforms using spike-sorting software provided by D. N. Hill, S. B. Mehta, and D. Kleinfeld ^5^. Single-units recorded on more than one session, as judged from the spike waveform and the firing pattern, were excluded from the analysis, but the results were not affected by the inclusion of all units. Recordings from outside the pPCx or CA1 were excluded from the analysis. We also excluded units with < 200 spikes in a given session. As in other cortical areas, neurons could be classified into two categories: wide-spiking (width > 0.2 ms) and narrow-spiking neurons (width < 0.2 ms). The width was determined as the time between a peak and a trough of the mean spike waveform. About 4 % (39) of recorded neurons fell into the category of narrow-spiking neurons. Overall, pPC neurons had spontaneous firing rates of 2.79 ± 3.74 Hz (mean ± S.D.). Spontaneous firing rates of wide-spiking neurons were significantly lower than those of narrow-spiking neurons (2.23 ± 2.49 Hz; 16.41 ± 3.43 Hz; mean ± S.D., respectively). Both categories of cells were included in the analyses but exclusion of narrow spike neurons did not affect the main conclusions.

#### Single neuron responsiveness and selectivity analysis

In order to obtain instantaneous firing rates (e.g. for Peri-event time histograms, PETHs), spike events were convolved with a Gaussian filter (S.D.: 25 ms). *<u>O</u> dors:* To identify odor responsive neurons (e.g. **Extended Data Fig 4d**), we compared the firing rate of a 1 second time window after odor onset to a 1 second time window immediately prior to odor onset using the Wilcoxon rank sum test, p<0.05. To identify neurons that were odor selective (e.g. **Fig. 2e**) we compared the firing rate during a 1 second time window after odor onset across the four odor stimuli using an ANOVA-test, p<0.05. *<u>Locations</u>*: Location selectivity of individual neurons was obtained from comparing firing rates in 20 x 20cm position bins centered around individual port locations on maze using ANOVA-test, p<0.05 (see ‘*Analysis of firing rate on maze’* section below).

#### Analysis of firing rate on maze

For firing rate heat map visualization (**Fig. 2, 3, Extended Data Fig. 3, Extended Data Fig 7b**), rat positions (tracked by LED on the microdrive) on the maze (1 meter x 1 meter) was divided into 4 x 4cm bins. Mean firing rate for each position bin normalized by occupancy was obtained and then smoothed with a gaussian kernel (S.D.: 2cm). Responses for each neuron used for location selectivity and correlation analysis was obtained by dividing the maze into 20 x 20 cm bins. The mean z-scored firing rate for the position bin centered on each port was taken as response at a particular location. Firing rate heat maps for individual odors (**Fig. 2 a-c**) were obtained by sorting trials by odor identity, and obtaining the firing rate meap map for all concatenated trials of the same odor. Firing rate heat maps for correct and error trials (**Extended Data Fig. 7b**) were generated by concatenating trials from -1 to 2 sec window around initiation and goal port entry in the entire session.

#### Population correlations (Fig 2, 3, Extended Data Fig. 4)

<u>Odor</u>: Population odor response vectors consisted of mean z-scored firing rate of neurons within a 1 second time window after odor onset. <u>Location:</u> Population location response vectors consisted of mean z-scored firing rate on the maze as described above in the ‘*Analysis of firing rate on maze’ section.* Pearson’s correlation coefficient was used for reported correlation coefficients. Qualitatively similar results were obtained using Spearman’s rank correlation coefficient.

#### Sparseness (Extended Data Fig. 4)

“Population sparseness” (*S_p_*) is defined for a given odor using the following formula, 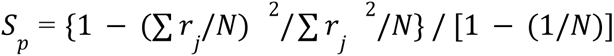, where N is the number of neurons and *r_j_* is the trial-averaged neural response (z-scored firing rate) of each neuron to the given odor <u>(Rolls and Tovee 1995; Willmore and Tolhurst 2001)</u>. “Lifetime sparseness” (*S_L_*) is calculated for each neuron using the above formula except *j* corresponds to each odor and N the total number of odors tested. Accordingly, *S_p_* quantifies sparseness of odor-evoked activity across a population for a given odor (0: uniform, 1: sparse). *S_L_* quantifies sparseness of odor responses (specificity) for a given neuron (0: uniform, 1: sparse). Population and lifetime sparseness for port locations were obtained similarly, using neural responses for port locations instead of odors.

#### Pseudopopulation decoding (Fig. 2g, k)

Subsets of neurons were sampled from across recording sessions. Population sizes were sampled at sqrt(2) increments up to the total number of available neurons leading to pseudopopulations of 2-725 (pPCx) and 2-91 (CA1). Neuronal responses were concatenated into a matrix, arranged so that each row contained responses to a particular (odor/location). Because different sessions contained different numbers of trials, a random subset of responses from each neuron was excluded so that the number of trials per category was consistent across neurons in the pseudopopulation. For each population size, the sampling procedure was repeated 10 times. Each neuron’s response was averaged over a 1-s window whose timing was chosen to maximize population decoding (see Extended Fig. 5; 0.25 to 1.25s post-odor onset for odor decoding; -0.5s to 0.5s peri-poke in for location decoding). Sniff-aligned odor decoding (Extended Fig. 5b) was performed by defining the first inhalation peak post-odor onset as t = 0. Each neuron’s response was normalized by taking the z-score across all trials. Trials were labeled 1-4 according to the odor sampled (for odor decoding) or port location (for location decoding), and arranged into a T (# trials)-element vector. Sessions with fewer than 400 trials per session was excluded, since it would lead to overall fewer usable trials in the entire pseudopopulation analysis. Multi-class classification was performed with the LIBLINEAR library^6^ for Matlab (https://www.csie.ntu.edu.tw/~cjlin/liblinear/), using either L1- or L2- regularized logistic regression with 5-fold cross-validation.

#### Population decoding

For per-session decoding, neuronal firing rates were arranged into a matrix of size T (# trials) x N (# neurons). Trials were labeled 1-4 according to the odor sampled (for odor decoding) or port location (for location decoding), and arranged into a T (# trials)-element vector. This matrix was normalized by taking the z-score of each column (so that each neuron’s firing rate was standardized). The vector of labels and matrix of firing rates were fed into a multi-class L1-regularized logistic regression (LIBLINEAR^6^, see previous section) using 5-fold cross validation. Chance-level classification was estimated by running the same classifier on shuffled labels. For decoder time courses (**Fig. 2h, l**), 200-ms non-overlapping windows were used. For pre-vs-post odor decoding (**Extended Data Fig. 6b, c**), time windows of 1-second before and after initial poke-in were used. Data sessions with fewer than 8 pPCx neurons were excluded because they had substantially fewer simultaneously recorded cells.

#### Population activity across epochs (Fig. 3a-c)

Initiation and goal epochs were defined as 1 second time windows centered around initiation and goal port poke-in times, respectively. Inter-trial interval (ITI) epoch was defined as the 4-6 second time window enforced in between trials. Occupancy of rat and firing rate heat maps for temporal epochs (Fig. 3a) were obtained by concatenating time periods of the same epoch.

#### Classification of location during correct and error trials

Each neuron’s average response over the interval [-.5s, .5s] peri-initiation port poke-in was z-scored across trials, and trained to classify port location as described in the ‘population decoding’ section. The trained decoder was used to classify the animal’s position during goal port poke-in for either correct or incorrect trials (**Fig. 3f**). For **Extended Data. Fig 7c**, the decoder was trained using only peri-initiation port poke-in for correct trials only. Sessions with fewer than 8 simultaneously recorded pPCx neurons were excluded.

#### Place field distribution along maze corridor (Fig. 3h)

Each neuron’s “place field” was defined as its mean firing rate in each of 10 equally-divided sections of each of the 4 arms of the plus-maze. Data was restricted to the 2-second period preceding poke-in, so that place fields reflected approach to port (and not departure). Neurons whose response along track never exceeded 2x the mean firing were excluded. Results were similar for other thresholds. Neurons were sorted by their place field locations along the arm containing peak response.

#### Arm identity decoding along maze corridor (Fig. 3i)

Data was restricted to the 1.5-second period preceding poke-in, which corresponds to the period where the animal is running up the maze arm (Fig. 3g). Neuronal firing rates were sampled at 5 Hz and z-scored, and the rat’s position was discretized into 5 equally-spaced bins along the arm length. 5 separate classifications were performed, one for each of the 5 bins, to predict which of the 4 arms the rat was occupying using either populations of simultaneously recorded pPCx or CA1 neurons. As with population decoding, classification was performed using LIBLINEAR^6^ with 5-fold cross-validation.

#### Noise correlation analysis (Extended Data Fig 8)

For each trial, each pPCx neuron’s activity was averaged over the 1-s period preceding poke-in. To remove potential signal correlations related to the animal’s position (odor responses did not apply to this pre-poke period), each neuron’s average response to the port was subtracted from the trial-specific firing rate. This was done separately for the first and second blocks of trials within a session, since the firing rates for some neurons drifted during the gap between trial blocks. Neurons were classified as significantly odor coding (O), significantly location coding (L), or neither. For pairs of neurons belonging to the same session, the pearson correlation coefficient was calculated and added to one of 6 sets corresponding to all possible pairings between O, N, and L. Note that O-O and L-L could contain the same neurons, since a neuron could be both odor- and location-coding. In contrast, the O-L group contained neurons that were exclusively odor or location coding, respectively. Results were similar when restricting O-O and L-L groups to be exclusively odor- or location-coding, respectively, or when increasing the sampling rate of neuronal firing (up to 16Hz, the maximum tested).

#### Identification of CA1-LFP

For each rat, the 96-electrode LFP data was represented as a 96-dimensional matrix sampled at 40 Hz. Each row of this matrix (representing a single electrode time series) was high-pass filtered using Matlab’s zero-phase ‘filtfilt’ function, and a 2Hz, 2nd order Butterworth filter. The resulting matrix was truncated to include only the time window [-1.5s, 2s] around the time of all trial-initiating poke-ins taking place within a session. For a given rat, the procedure was repeated for all sessions, and the resulting matrices were concatenated into a single matrix. In summary, this pre-processing captured each animal’s multi-electrode LFP during odor sampling and across sessions. SVD was applied to the resulting matrix, and the trial-averaged spectrograms (see below) of the top 10 components were visualized. To identify hippocampal theta, we looked for a component with power in the ∼8 Hz band^7^. For each rat, we picked the highest-variance component of this sort. Two additional lines of evidence suggested that we successfully identified theta: 1) theta power decreased around the time of poke-in, consistent with the observation that theta power decreases during periods of immobility (**Fig 4a**)^7^; and 2) the components showed strong coherence with identified hippocampal spiking activity at ∼8 Hz (see **Extended Data Fig 9d**, lower right panel) ^8^.

#### Spectral analysis

To compute power spectra and coherograms (**Fig. 4, Extended Data Fig 9**), the Chronux toolbox was used (http://chronux.org/) ^9^. Each neuron’s coherence with sniffing and CA1-LFP was calculated as a function of time and frequency using the function cohgramcpb, using Fs = 40 Hz, a sliding window of 1s with a stride of .2s, and taper set to [1 Hz, 1s, 1]. Similarly, the power spectrum for sniff and CA1-LFP was calculated using function specgram, with the same parameter settings. For this and subsequent analysis, data of 1 of 3 rats was excluded because it performed far fewer trials per session, leading to noisy estimates of coherence.

#### Single-cell decoding (Fig. 4b)

For each neuron, the firing rate was sampled at 2 Hz over the interval [-2s, 2s] with respect to initial poke-in (for location classification) and the interval [0s, 2s] with respect to odor onset (for odor classification). This was repeated across all trials, resulting in an R (# Trials) x T (# Timesteps) matrix. Each column of the matrix (i.e. each time point) was z-scored. Corresponding R-element vectors containing labels for poke-in location and odor identity was defined. For each, a linear multi-class decoder (using LIBLINEAR^6^, see above) was trained to classify odor/location based on the firing rate time series. This procedure was repeated 100 times for label-shuffled data to establish each neuron’s baseline classification performance, which deviated from 25% because of non-uniform trial sampling in sessions.

Finally, the mean chance-level performance was subtracted, and neurons whose classification accuracy exceeded the 95th percentile of the baseline classification accuracy were labeled as significant.

#### Relating single-cell decoding to coherence

*Correlation between decoding accuracy and coherence* (**Fig. 4c**): The coherograms of cells identified as significantly odor orlocationcoding (see above) were taken, and for each (time, frequency) value of the coherogram, the across-cell correlation between coherence and decoder was calculated. To identify (time, frequency) pairs with a significant association between decoding and coherence, the correlation was repeated 1,000x using shuffled decoder values, and (T,F) pairs whose value exceeded the 95th percentile were colored red (for sniff) and blue (for CA1-LFP).

#### Top ‘n’ cells analysis (Extended Data Fig. 9)

The top ‘n’ odor- and location- decoding cells were taken from Fig. 4a, and the difference of their mean coherograms was plotted. Significant (T,F) bins were identified as those whose values were significantly higher (lower) between the top ‘n’ location and top ‘n’ odor cells were colored blue (red), as indicated by a t-test (ttest2 in matlab).

## Notes

### Competing Interest Statement

The authors have declared no competing interest.

